# Protein coopted from a phage restriction system dictates orthogonal cell division plane selection in *Staphylococcus aureus*

**DOI:** 10.1101/2023.09.03.556088

**Authors:** Félix Ramos-León, Brandon R. Anjuwon-Foster, Vivek Anantharaman, Colby N. Ferreira, Amany M. Ibrahim, Chin-Hsien Tai, Dominique M. Missiakas, Jodi L. Camberg, L. Aravind, Kumaran S. Ramamurthi

## Abstract

The spherical bacterium *Staphylococcus aureus*, a leading cause of nosocomial infections, undergoes binary fission by dividing in two alternating orthogonal planes, but the mechanism by which *S. aureus* correctly selects the next cell division plane is not known. To identify cell division placement factors, we performed a chemical genetic screen that revealed a gene which we termed *pcdA*. We show that PcdA is a member of the McrB family of AAA+ NTPases that has undergone structural changes and a concomitant functional shift from a restriction enzyme subunit to an early cell division protein. PcdA directly interacts with the tubulin-like central divisome component FtsZ and localizes to future cell division sites before membrane invagination initiates. This parallels the action of another McrB family protein, CTTNBP2, which stabilizes microtubules in animals. We show that PcdA also interacts with the structural protein DivIVA and propose that the DivIVA/PcdA complex recruits unpolymerized FtsZ to assemble along the proper cell division plane. Deletion of *pcdA* conferred abnormal, non-orthogonal division plane selection, increased sensitivity to cell wall-targeting antibiotics, and reduced virulence in a murine infection model. Targeting PcdA could therefore highlight a treatment strategy for combatting antibiotic-resistant strains of *S. aureus*.

## INTRODUCTION

Bacterial cell division must be tightly regulated to ensure coordination between septum synthesis and faithful segregation of the genetic material. The central component of the division machinery in nearly all bacteria contains a tubulin homolog called FtsZ which assembles at mid-cell and directs the elaboration of the cell division septum ^1,2^. Correct placement of the cell division machinery has been extensively studied in rod-shaped model organisms such as *Escherichia coli* and *Bacillus subtilis* ^3–5^, but how bacteria that assume other shapes choose the correct division plane is poorly understood ^7^.

The spherical Gram-positive bacterium *Staphylococcus aureus* is a leading cause of bacteremia and nosocomial infections. As in most non-model bacteria, most components of the divisome have been identified by homology with those present in *E. coli* or *B. subtilis* ^8^. During the *S. aureus* cell cycle, cells replicate and segregate the nucleoid as FtsZ polymerizes into a ring at mid-cell. This is accompanied by a slight elongation of the cell, resulting in cells that are briefly ellipsoidal ^9^. After septum synthesis is completed, peptidoglycan hydrolysis is responsible for septum-splitting, a process that is extremely fast, resulting in two equally sized daughter cells ^10^. A distinctive feature of *S. aureus* cell division, first observed nearly fifty years ago, is that the two daughter cells divide in a cell division plane that is orthogonal to that of the parent cell ^6,11^, which results in *S. aureus* cells growing in grape-like clusters ^12^. The nucleoid occlusion protein, Noc, which prevents divisome assembly over the chromosome, has been suggested to be involved in this process, but deletion of the gene encoding Noc resulted in pleiotropic effects, which precluded a clear conclusion regarding the role of the nucleoid in positioning the cell division machinery in *S. aureus* ^13,14^. Thus, the mechanism by which the organism selects the correct cell division plane and the benefits, if any, of this unusual cell division pattern has been unclear.

To identify new cell division genes in *S. aureus*, we conducted a chemical genetic screen and identified PcdA, which is conserved in spherical Firmicutes that grow in clusters. Cells lacking PcdA failed to position the divisome orthogonal to the previous cell division plane. PcdA localized to future cell division sites in a nucleoid-independent manner, before FtsZ localized, and directly interacted with unpolymerized FtsZ to mark the correct cell division plane and assemble the cell division machinery at the proper localization. In the absence of PcdA, cells exhibited increased sensitivity to cell wall-targeting antibiotics and displayed decreased virulence in a mouse infection model, indicating that cell clustering by orthogonal cell division may represent a survival strategy against host immune defenses and environmental insults.

## RESULTS

### *pcdA* is required for orthogonal plane selection in *S. aureus*

To identify new cell division genes in *S. aureus*, we incubated individual strains in an ordered transposon library ^15,16^ in the absence or presence of a sublethal concentration of the FtsZ inhibitor PC190723 ^17^ and monitored growth over time. At this concentration of inhibitor, the wild type (WT) strain did not display significant growth reduction (Fig. 1a, black traces), but the cells were slightly larger compared to the absence of inhibitor (0.88 µm ± 0.24 µm, n=765 cells versus 1.2 µm ± 0.48 µm, n=840), indicating a slight cell division defect (Fig. 1b, c, h). However, one strain in the mutant library that exhibited a growth defect at this concentration of the inhibitor contained a transposon insertion in the *sausa300_2094* gene, which we renamed *pcdA* (<u>P</u>C190723-sensitive <u>c</u>ell <u>d</u>ivision <u>A</u>AA+ NTPase). To ensure that the growth defect was due to *pcdA* deletion, we constructed a marker-less deletion of *pcdA* and complemented it with a single copy of *pcdA* at an ectopic chromosomal locus. The absence of PcdA protein in the Δ*pcdA* strain was confirmed by immunoblot (Fig. S1a). Deletion of *pcdA* did not result in a growth defect, but in the presence of PC190723, the Δ*pcdA* strain displayed reduced growth compared to WT (Fig. 1a, pink traces), which was corrected upon complementation of the gene deletion in *trans* with a WT copy of *pcdA* (Fig. 1a, green traces). Additionally, the Δ*pcdA* strain displayed an altered morphology with a mean area of 1.2 µm^2^ ± 0.36 µm^2^, n=702 (Fig. 1d, h); in the presence of PC190723, this area increased to 1.5 µm^2^ ± 0.70 µm^2^, n=782 (Fig. 1e, h). These defects were complemented by expressing *pcdA* in *trans* (Fig. 1a, f, g, h). We next tested if deletion of *pcdA* resulted in a defect in orthogonal division plane selection by examining two consecutive cell division planes. First, we stained cell walls using fluorescently labeled wheat germ agglutinin (WGA) and washed away excess WGA. We then permitted one round of cell division, and stained cell membranes with the fluorescent dye FM4-64. Using fluorescence microscopy, we then measured the angle between the border of the WGA-stained region (which indicates the orientation of the previous cell division plane) and the septum labeled by FM4-64 (which indicates the current cell division plane) ^6^ (Fig. 1i-j’’). WT cells displayed a mean angle of 82.1° ± 9.89° (n=221) between each consecutive cell division plane, whereas Δ*pcdA* cells displayed a mean angle of 69.91° ± 24.6° (n=231) between each consecutive cell division plane, which included some cells that displayed cell division septa that were nearly parallel to the previous plane of division (Fig. 1k). This defect was largely restored upon complementation of *pcdA* in *tran* (81.5° ± 11.2° between each consecutive division plane, n= 203). Thus, deletion of *pcdA* disrupts proper selection of the division plane and renders cells hypersensitive to FtsZ inhibition.

**Figure 1.**
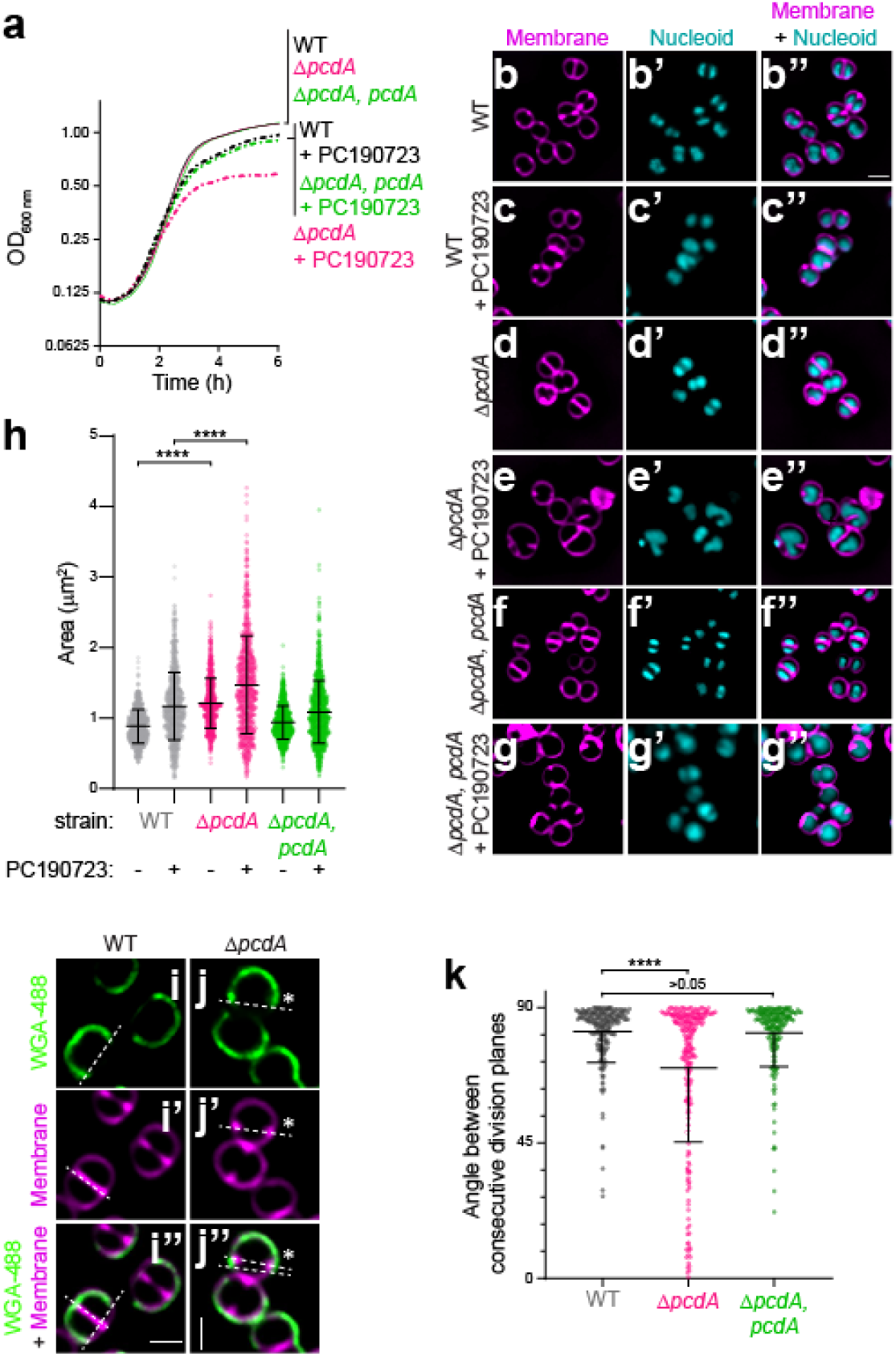
Deletion of *pcdA* results in a cell division defect. (a) Growth curves of WT (black), Δ*pcdA* (magenta), and Δ*pcdA* complemented at an ectopic chromosomal locus with *pcdA* (green) in rich media in the absence (solid lines) or presence (dashed lines) of 200 ng ml^-1^ FtsZ inhibitor PC190723. (b-g’’) Cell morphologies of (b-c’’) WT, (d-e’’) Δ*pcdA*, or (f-g’’) Δ*pcdA* complemented with *pcdA* in the (b-b’’, d-d’’, f-f’’) absence or (c-c’’, e-e’’, g-g’’) presence of PC190723 examined using fluorescence microscopy. b-g: membranes visualized with FM4-64 (magenta); b’-g’: nucleoid visualized using DAPI (cyan); b’’-g’’: overlay, membrane and nucleoid. Scale bar: 1 µm. (h) Cell sizes (calculated as area) of WT (gray), Δ*pcdA* (magenta), or Δ*pcdA* complemented with *pcdA* (green) strains in the presence or absence of PC190723 (n > 700 cells). Statistical analysis: one-way ANOVA; **** indicates p value < 0.001. (i-j’’) Representative fluorescence micrographs of (i) WT and (j) Δ*pcdA* stained with fluorescently labeled wheat germ agglutinin (WGA-488) and membrane dye FM4-64. WGA-488 was washed away and cells were allowed to divide for one round of cell division resulting in half cell staining. (i-j): fluorescence from WGA-488 (green); (i’-j’): membranes stained with FM4-64 (magenta); (i’’-j’’): overlay of WGA-488 and membranes. Division planes are indicated with dashed lines. Asterisk indicates a Δ*pcdA* cell with misplacement of the second division plane. Strains: JE2 and FRL60. (k) Angle between consecutive division planes in WT (gray), Δ*pcdA* (magenta), or Δ*pcdA* complemented with *pcdA* (green) strains. Bars indicate median; interquartile range indicated with whiskers. Strains: JE2, FRL60, and FRL62. Statistical analysis: Kruskal-Wallis; **** indicates < 0.0001.

### PcdA is a derived McrB family AAA+ NTPase with two N-terminal EVE domains

*pcdA* is present only in clump-forming coccoid Firmicutes related to *Staphylococcus* (e.g., *Mammalicoccus*, *Macrococcus*) but not in other lineages (Data Fig. S2a). Sequence-profile and profile-profile searches revealed that *pcdA* encodes a three-domain protein (Fig. 2a): the first two are tandem copies of the EVE domain (e-value: 10^-28^-10^-30^, PSI-BLAST iteration 2) ^18,19^ followed by a C-terminal AAA+-P-loop NTPase domain (HHpred probability: 99.86%, e-value: 2.6 x 10^-20^)^20^. EVE domains belong to the PUA (PseudoUridine synthase and Archaeosine transglycosylase)-like class of β-barrel domains that typically bind DNA or RNA with modified nucleobases ^19,21,22^. An examination of the multiple sequence alignment (MSA) and AlphaFold2-generated structural models of the PcdA EVE domains revealed that both possess the conserved cleft, which in other members of the PUA-like class are involved in the recognition of nucleic acids (Fig. 2b). An examination of the AAA+ NTPase domain revealed that it specifically belongs to the McrB family (Fig. S2a, Fig. 2c-d; HHpred e-value: 10^-18^-10^-20^, e.g., 6UT5, *Thermococcus gammatolerans* McrB), which primarily includes the NTPase subunits of restriction systems that target DNA of phages with base modifications such as 5-methylcytosine and 5-hydroxymethylcytosine. The McrB family belongs to a clade of AAA+ domains that is typified by two diagnostic β-hairpin inserts: one in the middle of helix-2 of the core P-loop domain and the other just prior to sensor-1 (Fig. 2d). Our MSA and structural models show that PcdA has both these β-hairpins (Fig. 2c).

**Figure 2.**
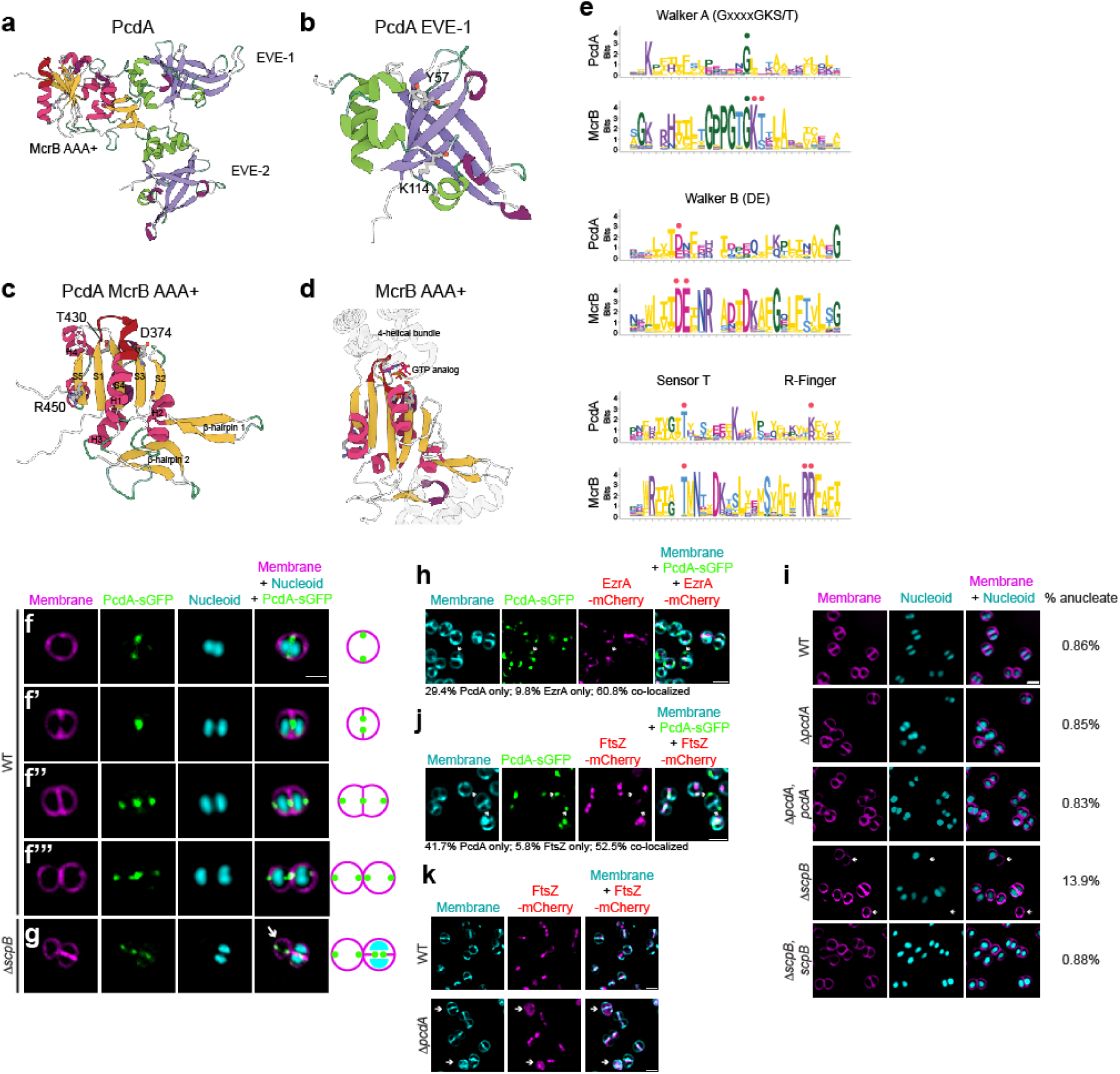
PcdA is an early cell division protein that belongs to the AAA+ family of NTPases. (a-d) Cartoon representations of the predicted AF2-generated structures of (a) full length PcdA, (b) EVE domain 1 of PcdA, and (c) McrB AAA+ domain of PcdA. (d) Crystal structure of *Thermococcus gammatolerans* McrB (PDB: 6UT3). Select residues mentioned in the text are marked. (e) Sequence logo displaying conservation of amino residues in indicated motifs of the AAA+ domains of PcdA and McrB orthologs. The height of each residue is scaled as per the bitscore of conservation in the MSA, measured using Shannon entropy. Red dots: key active site residues; green dots: other conserved sites. (f-g) Subcellular localization of PcdA-sGFP in (f) pre-divisional, (f’) nascently dividing, (f’’) nearly completely divided, or (f’’’) completely divided *S. aureus* cell, or (g) in a representative anucleate Δ*scpB* mutant *S. aureus* cell. Arrow indicates an anucleate cell. First column: membranes visualized using FM4-64 (magenta); second column: PcdA-sGFP (green); third column: nucleoid visualized using DAPI (cyan); fourth column: overlay of membrane, nucleoid, and PcdA-sGFP. Schematic representation of PcdA localization (green: PcdA; magenta: membrane) shown to the right. Strains used: FRL28 and FRL68. (h) Colocalization of PcdA-sGFP and EzrA-mCherry. Panels from left to right: membranes visualized using TMA-DPH (cyan); PcdA-sGFP (green); EzrA-mCherry (magenta); overlay of membranes, PcdA-sGFP, and EzrA-mCherry. Arrow indicates a representative PcdA-sGFP focus without colocalization of EzrA-mCherry. Localization frequencies of PcdA-sGFP or EzrA-mCherry alone, or colocalization of both proteins are indicated below. Strain FRL109. (i) Analysis of presence of anucleate cells in the JE2 wild type strain (WT), Δ*pcdA*, complemented Δ*pcdA*, Δ*scpB*, and complemented Δ*scpB*. First column shows membrane in magenta stained with FM4-64; second column shows nucleoid in cyan stained with DAPI; and third column shows the overlay of the two previous images. Percentage of anucleate cells for each strain is indicated in the fourth column (n > 1000 cells). Strains: FRL60, FRL62, NE1085, and FRL12. (j) Co-localization of PcdA-sGFP and FtsZ-mCherry. Panels from left to right: membranes visualized using TMA-DPH (cyan); PcdA-sGFP (green); FtsZ-mCherry (magenta); overlay of membranes, PcdA-sGFP, and FtsZ-mCherry. Arrow indicates a representative PcdA-sGFP focus without colocalization of FtsZ-mCherry. Localization frequencies of PcdA-sGFP or FtsZ-mCherry alone, or colocalization of both proteins are indicated below. Strain FRL117. (k) Localization of FtsZ-mCherry in WT and Δ*pcdA*. First column: membranes visualized with TMA-DPH (cyan); second column: FtsZ-mCherry (magenta); third column: overlay of membrane and FtsZ-mCherry. Scale bars, 1 μm. Arrows indicate cells where FtsZ-mCherry signal is soluble and not forming rings. Strains: FRL115 and FRL116.

Members of the McrB family are characterized by either of two architectural themes. Most commonly, the AAA+ NTPase domain is fused to one or more N-terminal “reader” domains that specifically recognize base modifications in particular sequences of invading phage DNA to discriminate it from unmodified host DNA ^19,22^. Alternatively, especially in multicellular bacteria, the AAA+ NTPase domain is coupled to “co-effector” domains or “effector-associated” domains (EAD)/Death-like superfamily domains that are predicted to either directly or indirectly trigger (via recruitment of another effector through homotypic interactions) a suicide response upon failure of restriction due to phage counterattack ^23^. While the modified DNA reader domains belong to several structurally diverse folds, one of the most common throughout the McrB family are the EVE domains. Thus, the above observations firmly place PcdA within the classical McrB family of AAA+ NTPases. Indeed, consistent with our sequence searches, a phylogenetic tree recovers PcdA as a divergent branch of an McrB subclade that is enriched in Firmicutes (Bacillota; Fig. S2a).

Despite the conservation, PcdA differs from classic McrBs in multiple ways. In structural terms, PcdA has lost the C-terminal four helical bundle that is characteristic of AAA+ NTPases (Fig. 2c-d). Keeping with this structural degeneration, the Walker A motif is largely degraded, the Walker B motif has lost the glutamate downstream of the conserved Mg^2+^-chelating aspartate, and one of two successive arginine fingers occurring at the helix-4—strand-5 junction (an McrB family-specific feature) is lost (Fig. 2e). Finally, most members of the McrB family involved in modified DNA restriction occur in an operon with a second gene coding for a restriction enzyme (typically McrC). The restriction subunit has a characteristic two-domain architecture with a restriction-endonuclease fold domain linked to an N-terminal McrC-NTD that activates GTP hydrolysis by the McrB AAA+ domain ^24–26^. Such a linked restriction subunit with an McrC-NTD is absent in PcdA. Together, these observations suggest that PcdA, while emerging from a bona fide ancestral McrB protein, has likely undergone a major functional shift. Importantly, while it might still bind a nucleotide (due to the intact Mg^2+^-chelating residue) and form a multimer (Fig. S2b-e), it is predicted to exhibit weak, if any, NTPase activity.

### PcdA is an early cell division protein

To test if PcdA is a divisome component, we analyzed the subcellular localization of PcdA fused to superfolder GFP (PcdA-sGFP) using fluorescence microscopy (Fig. 2f-f’’’, larger field of view in Fig. S1c). In cells that had not yet initiated septation and the nucleoid had begun to segregate, PcdA-sGFP formed a ring at mid-cell (as evidenced by two puncta when viewed at an intermediate focal plane; Fig. 2f). In cells that were actively constricting, the PcdA-sGFP ring localized at the leading edge of the division septum, suggesting that it was co-constricting with the cell division machinery (Fig. 2f’). Once cells had completed septation but had not yet split, a population of PcdA-sGFP localized as a ring in each daughter cell at an orthogonal plane corresponding to the next site of cell division, while a second population of PcdA-sGFP remained near the site of constriction of the nascently formed division septum (Fig. 2f’’). In newly split cells that remained attached but had not yet initiated chromosome segregation, each daughter cell harbored a PcdA-sGFP ring coincident with the next plane of cell division (Fig. 2f’’’). The presence of EVE domains in PcdA, which are typically implicated in interactions with modified nucleic acids, made us wonder if the chromosome could participate in PcdA localization. To address this question, we examined PcdA-sGFP localization in mutant cells (Δ*scpB*), which are defective for chromosome segregation and therefore generate an increased number of anucleate cells. In anucleate cells, PcdA-sGFP localized as a ring at mid-cell at a similar plane as its sister cell that contained nucleoids, indicating that the nucleoid is not required for proper localization of PcdA (Fig. 2g).

The redeployment of PcdA, from mid-cell to the future division planes, in daughter cells that had not yet split suggested that PcdA arrives very early at the division site. To measure this, we examined the co-localization of PcdA-sGFP with EzrA-mCherry, an early cell division protein that is a scaffold for the assembly of the division machinery ^27^. In 60.8% of cells that elaborated complete septa, PcdA-sGFP co-localized with EzrA-mCherry (Fig. 2h). However, in 29.4% of cells, PcdA-sGFP localized at a division site without a corresponding EzrA-mCherry signal, while only 9.8% of cells (n=51 cells) harbored an EzrA-mCherry signal without a co-localized PcdA-sGFP signal, suggesting that PcdA is an early cell division protein that arrives at the division site earlier than EzrA. Finally, since cell division and chromosome segregation are often tightly linked processes in bacteria, we examined if PcdA plays a role in segregating chromosomes. Deletion of *scpB*, which is part of the SMC complex, resulted in 13.9% (n = 3494) of cells being devoid of nucleoid, whereas far fewer WT (0.86%, n = 5059) or Δ*pcdA* (0.85%, n = 1656) cells were anucleate (Fig. 2i, Fig. S1d). Together, the results indicate that PcdA is an early cell division protein that localizes to new division sites in a nucleoid-independent manner and that it likely controls cell division plane selection without influencing chromosome segregation.

To understand the relationship between PcdA and FtsZ, we examined the co-localization of PcdA-sGFP and FtsZ-mCherry. To prevent pleiotropic effects of FtsZ overproduction, FtsZ-mCherry was expressed under a Cd^2+^-inducible promoter, and its production was induced using 0.5 µm CdCl_2_ for 30 min while cells were actively growing in mid-exponential phase. In actively dividing cells, both proteins co-localized in 52.5% of cells (n = 120; Fig. 2j). However, in 41.7% of these cells, we observed only PcdA-sGFP localization without a corresponding FtsZ-mCherry signal, whereas 5.8% of cells displayed only FtsZ-mCherry signal, suggesting that PcdA localizes to the division site before FtsZ. We next analyzed the localization of FtsZ-mCherry in the absence of PcdA. In Δ*pcdA* cells, FtsZ-mCherry appeared as a soluble signal in the cytosol in 25.9% of cells (n = 1011), compared to only 4.2% in otherwise WT cells (n = 883; Fig. 2k), suggesting that FtsZ failed to efficiently polymerize as a ring in the absence of PcdA. Given the dependence of FtsZ on PcdA for localization and polymerization, the data thus far were consistent with a model in which PcdA acts as a positive regulator for Z-ring placement.

### PcdA possesses NTPase activity despite extensive sequence divergence

We next tested the *in vivo* requirements of the different domains of PcdA using site-directed mutagenesis of key residues, guided by sequence alignment and structural predictions.

Alanine substitutions were carried out in two conserved residues predicted to form part of the classical nucleic-acid-binding interface in each of the EVE domains: Y57A and K114A in EVE-1, and Y187A and K253A in the EVE-2 domain (Fig. 2b). Alanine substitutions were also made to the following residues of the McrB AAA+ NTPase domain: K313A associated with the remnant of the Walker A (“A*”), D374A (the Mg^2+^-chelating residue of the degenerate Walker B motif; “B*”),

T430A (intact sensor threonine, “T*”), and R450A (putative arginine finger, “R*”) (Fig. 2c). These PcdA variants were expressed in the Δ*pcdA* strain, and their expression and stability were confirmed by immunoblot blot (Fig. S3a). Functionality of these variants was then analyzed by the ability of each variant to complement the cell size defect of the Δ*pcdA* strain (Fig. 3a). Disruption of conserved residues in EVE-1 or EVE-2 resulted in increased cellular area, similar to the deletion of *pcdA*. Despite the defective nature of the Walker A motif, substituting the Lys residue resulted in a 21% increase in mean cellular area compared to WT. Similarly, despite the absence of the conserved Glu in the defective Walker B motif (Fig. 2a), substituting the conserved Asp increased mean cellular area to that observed with cells harboring a *pcdA* deletion (Fig. 3a). Finally, cells harboring the T* and R* variants of PcdA showed 24.1% and 33.0% increases in cellular area compared to WT (Fig. 3a). Thus, disrupting residues in both the EVE and AAA+ NTPase domain affect PcdA function in vivo.

**Figure 3.**
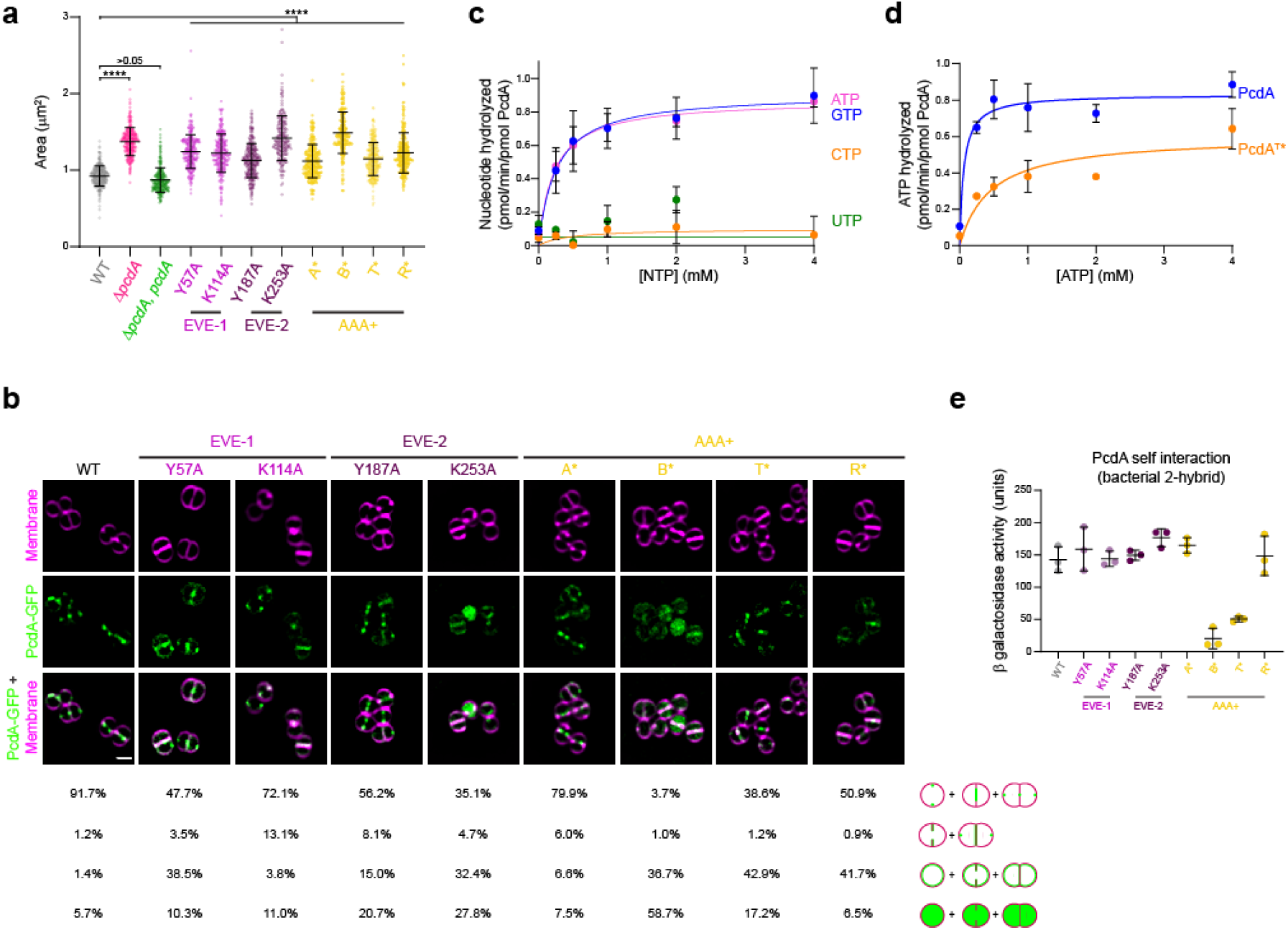
PcdA ATPase activity is necessary for its function and localization. (a) Cellular area (μm^2^) of the indicated strains (n > 300). Bars indicate the median with interquartile range. Strains: JE2, FRL60, FRL14, FRL34 – 41. Statistical analysis: one-way ANOVA; **** indicates p < 0.0001. (b) Localization of PcdA-sGFP and variants. First row shows membranes in magenta stained with FM4-64; second row shows PcdA-sGFP in green; and third row shows overlay of the two previous images. Below, localization of PcdA was quantified as correct localization (constricting ring as septation occurs; fourth row), non-constricting ring (fifth row), mis-localized all over the membrane (sixth row), or soluble localization (seventh row). Percentage of each type of localization are indicated for each protein variant (n > 300). Strains: FRL28, FRL44 – 51. (c) Nucleotide hydrolysis rate of PcdA. 2.5 μM PcdA was incubated with increasing concentration of ATP, GTP, CTP, or UTP (0, 0.25, 0.5, 1, 2, and 4 mM). NTP hydrolysis was quantified by the release of inorganic phosphate. Errors: S.D. (n = 5 independent experiments). (d) ATP hydrolysis rate for wild type PcdA and PcdA^T*^. 2.5 μM each variant was incubated with increasing concentrations of ATP (0, 0.25, 0.5, 1, 2, and 4 mM). Errors: S.D. (n = 3 independent experiments). (e) Multimerization of PcdA and mutated variants studied by bacterial two hybrid. The interaction of the proteins produced by the T18 and T25 plasmids cloned in a *cyaA* deficient *E. coli* was measured as β-galactosidase activity in liquid cultures. The protein variant fused to the N-terminus of T18 and T25 is indicated. A pair of non-fused T18 or T25 together with their corresponding fusion protein was used as a control and the resulting activity from the control was used to subtract to the tested interaction. Bars represent mean; whiskers represent S.D. (n = 3 independent experiments).

Next, we examined the subcellular localization of the PcdA variants (Fig. 3b). Whereas WT PcdA-sGFP localized at the septum as a constricting ring in 91.7% (n = 428) of cells, disrupting either EVE domain increased cytosolic distribution of PcdA, or abrogated PcdA redeployment to the new division plane, with PcdA instead mis-localizing uniformly to the membrane during and after septation (Fig. 3b, columns 2-5). Notably, substituting Lys in EVE-1 disrupted the ability of this variant in migrating with the leading edge of the constricting membrane (Fig. 3b, column 3). In the AAA+ domain, the PcdA^A*^ variant showed only a modest defect in localization compared to WT but disrupting the defective Walker B motif nearly abolished localization to the divisome or to the new plane of cell division, with most of PcdA^B*^-sGFP residing in the cytosol or indiscriminately in the membrane (Fig. 3b, columns 6-7). Disrupting the sensor T or arginine finger also reduced proper localization of PcdA mostly by increased indiscriminate mis-localization of these variants in the membrane (Fig. 3b, columns 8-9). The data therefore indicate that, despite the disrupted nature of signature motifs in the P-loop of the AAA+ domain, NTP binding and possibly weak NTP hydrolysis is critical for PcdA function.

Since the *in vivo* analyses suggested a key role for the AAA+ domain in PcdA function, we sought to test the NTPase activity of the protein in vitro. We therefore purified untagged recombinant PcdA and measured its ability to hydrolyze various nucleotides. Incubation of increasing concentrations of either ATP or GTP with PcdA produced saturation curves that revealed substrate turnover rates (*k*_cat_) of 0.93 ± 0.09 pmol ATP min^-1^ pmol^-1^ PcdA and 0.91± 0.12 pmol GTP min^-1^ pmol^-1^ PcdA, suggesting that PcdA does not distinguish between these nucleotides (Fig. 3c). In contrast, we did not observe significant hydrolysis of CTP or UTP. As a control for specificity of this relatively low level of hydrolysis, we observed that the purified T* variant of PcdA displayed reduced turnover rate (0.60 ± 0.08 pmol ATP min^-1^ pmol^-1^ PcdA^T*^) and reduced catalytic efficiency (*k*_cat_/*K*_m_) for ATP hydrolysis (1.33 min^−1^ mM^−1^) compared to WT PcdA (13.7 min^−1^ mM^−1^; Fig. 3d).

Disruptions to the nucleotide binding pocket of AAA+ proteins can affect their ability to multimerize ^28^. We therefore tested the ability of PcdA variants to self-interact using a bacterial two hybrid assay^29^ by expressing PcdA or variants fused to adenylate cyclase subunit T18 and T25 in *E. coli* and analyzing self-interaction by measuring β-galactosidase activity. WT PcdA displayed robust self-interaction in this assay (Fig. 3e), as did variants that disrupted either EVE domain. In the AAA+ domain, although disrupting the already defective Walker A motif or the putative arginine finger did not affect PcdA self-interaction, disrupting either the defective Walker B motif or the sensor threonine abrogated PcdA self-interaction. Taken together, despite a very low NTPase activity (consistent with a degenerate P-loop and absence of the Walker B general base glutamate), we conclude that nucleotide hydrolysis nonetheless is required for multimerization, proper subcellular localization, and function of PcdA.

### PcdA interacts directly with FtsZ and DivIVA

PcdA localization at mid-cell along with the divisome led us to hypothesize that PcdA could interact directly with FtsZ. To investigate this, we purified recombinant untagged *S. aureus* FtsZ and PcdA and performed co-sedimentation assays in the absence and presence of the non-hydrolysable GTP analog GMPCPP (Fig. 4a). In the presence, but not in the absence, of GMPCPP, nearly half of FtsZ was found in the pellet, indicating nucleotide-dependent FtsZ polymerization (Fig. 4a, lanes 1-4). In contrast, the presence of GMPCPP did not increase the amount of PcdA in the pellet, suggesting that PcdA alone does not polymerize with GMPCPP (Fig. 4a, lanes 5-8). However, when FtsZ and PcdA were incubated together in the presence of GMPCPP, the majority of PcdA was detected in the pellet (along with over half of the FtsZ in the reaction), indicating that PcdA can directly interact with polymerized FtsZ (Fig. 4a, lanes 9-12). To test if PcdA can bind to unpolymerized FtsZ, we performed filter retention assays in which we incubated FtsZ and PcdA in the absence of GTP (to ensure that FtsZ would not polymerize), but in the presence of ATP, which is hydrolyzed only by PcdA, and centrifuged the samples through a filter that would retain protein complexes larger than 100 kDa (Fig. 4b). In the absence of ATP, PcdA alone, FtsZ alone, or FtsZ combined with PcdA did not show appreciable retention in the filter, indicating that they did not form a complex under these conditions (Fig. 4b, lanes 1-6). In the presence of ATP, neither PcdA alone nor FtsZ alone was retained (Fig. 4b, lanes 7-10), but when combined, PcdA and FtsZ were both retained on the filter (Fig. 4b, lanes 11-12), indicating that PcdA and FtsZ interact in the absence of GTP in an ATP-dependent manner.

**Figure 4.**
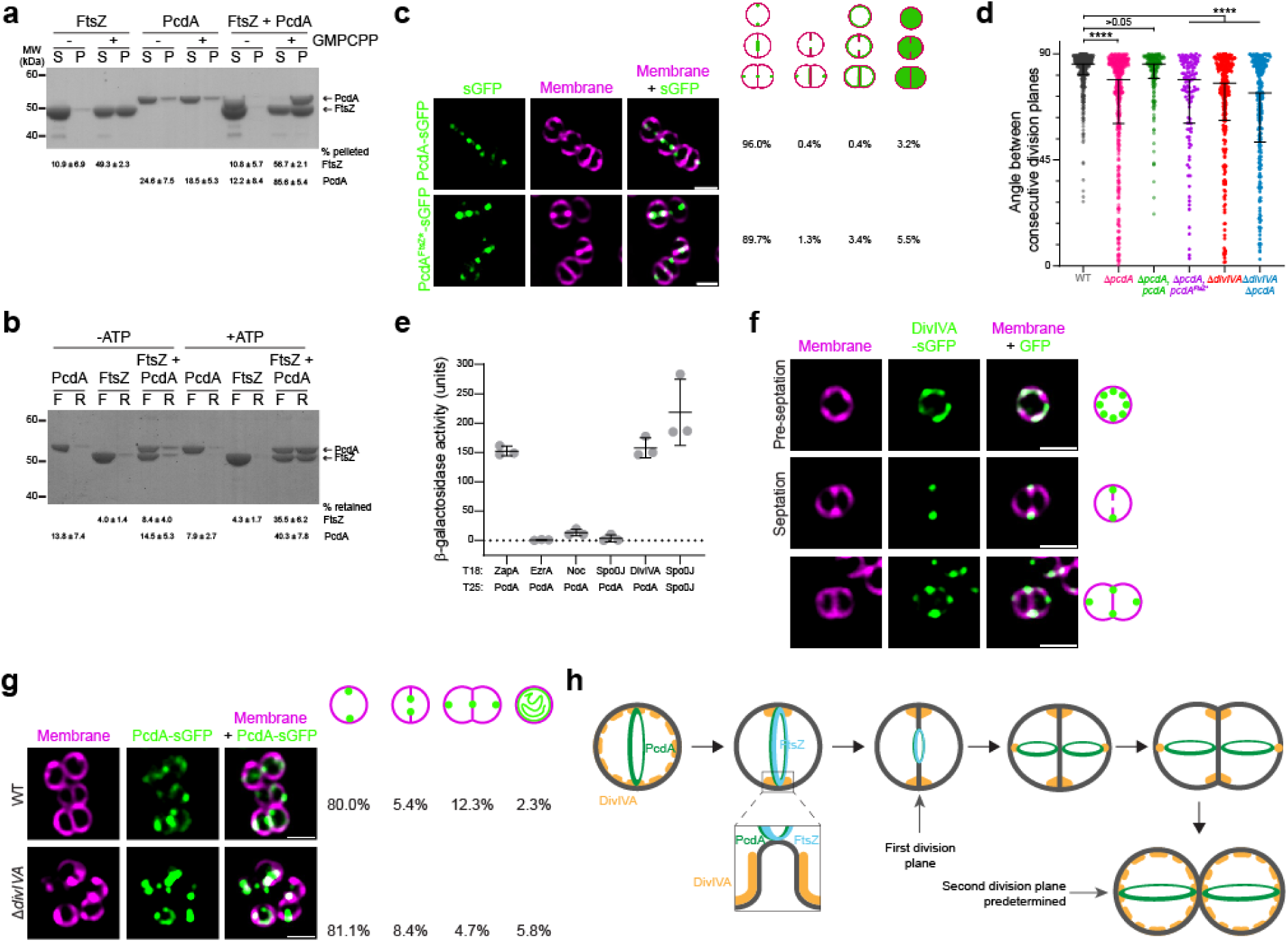
PcdA interacts directly with FtsZ and DivIVA. (a) Co-sedimentation of PcdA with polymerized FtsZ *in vitro*. 30 μM FtsZ was incubated in the presence or absence of 2 mM GMPCPP and 5 μM PcdA as indicated. FtsZ polymers were collected by high-speed ultracentrifugation. Presence of proteins in the supernatant (S) or pellet (P) was analyzed by SDS-PAGE and Coomassie staining. A representative image of three independent experiments is shown. The mean and standard deviation of the percentage of total FtsZ and PcdA in the pellet fraction is indicated below. (b) Interaction of PcdA with unpolymerized FtsZ depends on the presence of ATP. 30 μM FtsZ and/or 5 μM PcdA were incubated in the absence or presence of ATP as indicated. Protein mixture was then applied to a 100 kDa filter and centrifuged. Flowthrough (F) and resuspended retained protein (R) was analyzed by SDS-PAGE and Coomassie staining. The mean and standard deviation of the percentage of total FtsZ and PcdA in the retained fraction is indicated below. (c) Localization of PcdA and a variant with mutations in the predicted FtsZ-interacting residues. First column: fluorescence from sGFP variant (green); second column: membranes stained with FM4-64 (magenta); third column: overlay sGFP and membrane. To the right, localization of PcdA was quantified as correct localization (constricting ring as septation occurs; fourth column), non-constricting ring (fifth column), mis-localized all over the membrane (sixth column) or soluble localization (seventh column). Percentage of each type of localization are indicated for each protein variant (n > 400). Strains: FRL103 and FRL84. (d) Angle between consecutive division planes using cell wall staining with WGA-488 and membrane dye FM4-64 ^6^. Bars represent median values with interquartile range (n > 100 cells). Strains: JE2, FRL60, FRL62, FRL83, FRL96, and FRL98. Statistical analysis: Kruskal-Wallis; **** indicates p < 0.0001. (e) Interaction of PcdA with other cell division proteins by bacterial two-hybrid. PcdA was fused to the N-terminus of T18 subunit and expressed together with the indicated staphylococcal cell division protein fused to the N-terminus of the T25 subunit. Interaction between PcdA-T18 and the T25-fused cell division protein led to detection of β-galactosidase activity. Known self-interaction between Spo0J was used as positive control. (f) Subcellular localization of DivIVA-sGFP in a representative (first row) pre-divisional, (second row) nascently dividing, or (third row) nearly completely divided *S. aureus* cell. First column: membranes visualized with FM4-64 (magenta); second column: DivIVA-sGFP (green); third column: overlay of membrane and DivIVA-sGFP. Scale bar, 1 μm. Strain FRL113. (g) Localization of PcdA-sGFP in (top row) WT and (bottom row) Δ*divIVA*. First column: membranes visualized with FM-4-64 (magenta); second column: PcdA-sGFP (green); third column: overlay of membrane and PcdA-GFP. Percentage of each type of indicated localization pattern are shown to right (n > 300). Scale bars, 1 μm. Strains: FRL103 and FRL97. (h) Model for cell division plane selection in *S. aureus*. In pre-divisional cells, DivIVA (yellow) localizes indiscriminately in the membrane and PcdA (green) localizes to the future cell division plane, where it recruits FtsZ (blue). As the membrane invaginates, DivIVA localizes to the base of the nascent septum and PcdA follows the leading edge of the constricting septum. As cytokinesis completes, a population of DivIVA deploys to the poles of the slightly elongated cell, where it forms patches and recruits PcdA, which begins assembling as a ring defining the next division plane, orthogonal to the previous plane.

Next, we modeled the interaction between PcdA and FtsZ using AlphaFold-Multimer ^30^ (Fig. S3b). The model predicted that three residues in the EVE-1 domain of PcdA (R16, E31, and Q60) reside on a surface that could interact with the N-terminal domain of FtsZ. We disrupted this interaction surface by substituting each residue with Ala, complementing the Δ*pcdA* strain with this putative FtsZ interaction-deficient allele of *pcdA* (“*pcdA*^FtsZ^*”), and examined the angle between consecutive division planes in cells producing PcdA^FtsZ^*. The subcellular localization of angle between consecutive division planes revealed that *pcdA*^FtsZ^* was unable to complement the division defect of the Δ*pcdA* strain (Fig. 4d, lanes 1-4). Thus, reducing the interaction between PcdA and FtsZ by disrupting the putative interaction surface between the two proteins specifically disrupted division plane selection, but not PcdA localization. The results are therefore consistent with a model in which PcdA localizes to the correct cell division plane and subsequently directly recruits FtsZ to that site.

To understand how PcdA localizes correctly, we tested the interaction of PcdA with known cell division proteins using the bacterial two-hybrid assay ^31^. As a control, the partitioning protein Spo0J showed robust self-interaction in this assay ^32^ (Fig. 4e, lane 6). We did not detect appreciable interaction between PcdA and EzrA, Noc, or Spo0J ^13,14,27^ (Fig. 4e, lanes 2-4). However, we did detect interaction between PcdA and the early cell division protein ZapA ^33^ (Fig. 4e, lane 1) and the multifunctional coiled-coil structural protein DivIVA (Fig. 4e, lane 5) that is widely conserved in several bacterial lineages ^34^. In *B. subtilis*, DivIVA mediates the localization of the “Min” regulators of cell division ^35,36^, but the function of this protein in *S. aureus* has been mysterious ^37,38^. We therefore deleted *divIVA* in the Δ*pcdA* background and analyzed the phenotype of this strain. The Δ*pcdA* Δ*divIVA* strain displayed a similar cellular area as the Δ*pcdA* strain (Fig. S3c) and showed a similar defect in the selection of the orthogonal cell division plane (Fig. 4c, lane 6). Complementation of *pcdA* in the Δ*pcdA* Δ*divIVA* strain resulted in a cellular area similar to that of WT cells (Fig. S3c), but still resulted in a defect in orthogonal plane selection (Fig. 4c, lane 5), suggesting that PcdA and DivIVA work together in cell division plane selection, but that PcdA plays an additional role during the cell division process itself. Given the role of DivIVA in positioning proteins in other systems, the results are consistent with a model in which DivIVA recruits PcdA to the correct cell division site, which, in turn, recruits unpolymerized FtsZ to begin assembly of the divisome at the correct cell division plane.

To test the relationship between PcdA and DivIVA, we examined DivIVA-sGFP localization in the absence of PcdA. In WT cells, DivIVA-sGFP localization depended on the stage of the cell cycle (Fig. 4f, Fig. S3d). In cells that had not yet started septation (Fig. 4f, top row), DivIVA-sGFP displayed a patchy localization pattern along the membrane. In cells that had started septation (Fig. 4f, middle row), DivIVA-sGFP localized near the base of the division septum, similar to the pattern observed in *B. subtilis* ^36^, but unlike what we observed for PcdA-sGFP (which followed the leading edge of the division septum). In cells that had nearly completed cytokinesis but had not yet separated into two daughter cells (Fig. 4f, bottom row), an additional population of DivIVA-sGFP localized to the future cell division plane, forming foci that resembled those formed by PcdA. In the Δ*pcdA* strain, localization of DivIVA-sGFP was similar to that of WT, with only an increase in those cells displaying DivIVA-sGFP at the division site during septation (6.8% cells in WT, n = 413; 16.0% cells in Δ*pcdA*, n = 481; Fig. S3d). In contrast, deletion of *divIVA* reduced the frequency of cells in which PcdA-sGFP redeployed to the new cell division plane (Fig. 4g; 12.3% in WT; 4.7% in Δ*divIVA*). The results therefore suggest that DivIVA localization is not dependent on PcdA, but redeployment of PcdA to the next cell division plane depends on DivIVA.

### Deletion of *pcdA* reduces virulence and increases sensitivity to cell wall-targeting antibiotics

Since deletion of *pcdA* permitted growth in rich medium, we sought to understand the benefit of orthogonal cell division plane selection in *S. aureus* by testing the virulence of the Δ*pcdA* strain in a murine intravenous infection model. Following inoculation in the bloodstream, *S. aureus* disseminates to organs, such as kidneys, where they form abscesses within 4-5 days and persist for over 15 days ^39,40^. We therefore infected mice with WT or Δ*pcdA* cells, harvested kidneys on days 5 or 15 post-infection, and enumerated abscesses on the surface of the kidneys. Although the number of abscesses initially formed were equivalent in both infections, on day 15 the Δ*pcdA* strain displayed a 2.8-fold reduction in abscess formation (Fig. 5a). Histological analysis of the kidneys isolated at day 5 post-infection of both WT and Δ*pcdA* strains displayed the classical architecture of abscess lesions with prominent infiltration of immune cells surrounded by a clear layer of fibrin demarcating infected and healthy tissues (Fig. 5b-c’). However, enumeration of the presence or absence of bacteria at the center of these lesions revealed that, while infection with the WT strain resulted in 66% ± 13% of total lesions that contained bacteria, only 37% ± 11% of lesions resulting from infection with the Δ*pcdA* strain contained bacteria (Fig. 5d), indicating that deletion of *pcdA* likely increases the susceptibility of the bacterium to clearance by the host.

**Figure 5.**
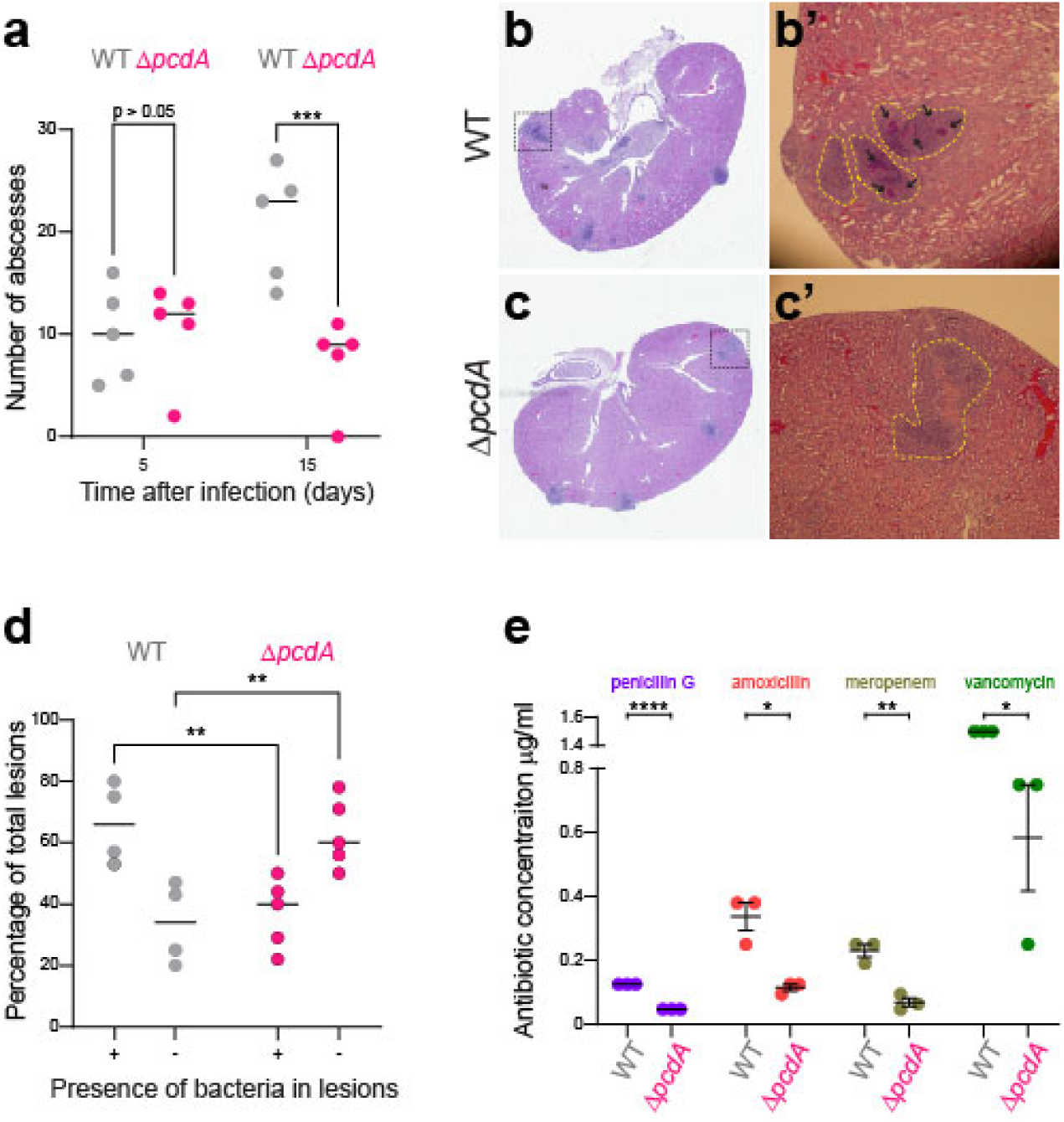
Deletion of *pcdA* impairs virulence and leads to increased sensitive to cell wall-targeting antibiotics. (a) Quantification of abscesses 5-and 15-days post infection. Mice were inoculated with WT or Δ*pcdA* strain. Mice were sacrificed after 5 or 15 days and the number of abscesses present in the kidneys was determined. Number is plotted as the mean from 5 animals per group. (b-c) Histopathology of kidneys of mice inoculated with JE2 wild type (b-b’) and Δ*pcdA* (c-c’). Pathological section was stained with hematoxylin and eosin (H&E). B’ and C’ show a close-up image of the lesions that are traced by yellow dotted line. Black arrows point to *S. aureus* cells growing inside the lesion. (d) Percentage of lesions with or without bacteria in kidneys of mice inoculated with JE2 wild type or Δ*pcdA*. Statistical analysis: two-way Anova; ** indicates p < 0.01; *** indicates p < 0.001. (e) Minimal inhibitory concentrations (MIC) for JE2 wild type and Δ*pcdA* for different antibiotics that target the cell wall. MIC was determined in lawns of bacteria using MIC strips for the indicated antibiotic. Strains: JE2 and FRL60.

We next tested the susceptibility of the Δ*pcdA* to various clinically relevant antibiotics. When challenged with antibiotics that target the cell wall, the Δ*pcdA* strain displayed reduced minimal inhibitory concentrations (MIC) for penicillin (2.7-fold relative to WT), amoxicillin (3-fold), meropenem (3.4-fold), and vancomycin (2.6-fold) (Fig. 5e). In contrast, the susceptibility of the Δ*pcdA* strain to antibiotics that target other cellular processes such as membrane integrity, DNA metabolism, or protein synthesis was largely unchanged (Fig. S4). Taken together, we conclude that disrupting the selection of the proper cell division plane leads to a virulence defect and increased susceptibility to cell wall targeting antibiotics that are commonly used to combat *S. aureus* infections.

## DISCUSSION

Nearly all genes involved in cell division in *Staphylococcus aureus* were identified by homology with those present in non-spherical bacteria such as *Bacillus subtilis* and *Escherichia coli* ^8^, and even novel cell division genes reported first in *S. aureus* are typically conserved in non-spherical cells ^41^. Although study of these conserved genes has led to a better understanding of cell division in *S. aureus*, important aspects are still poorly understood due to the absence of homologs of key components such as the Min system, involved in controlling Z-ring placement in other bacteria ^36,42^. While factors such as nucleoid occlusion factor Noc have been proposed to play a role in Z-ring positioning, deletion of such genes usually have pleiotropic effects, so it is not always clear whether Z-ring positioning is the primary function of that gene ^13^. Here we showed that the McrB family AAA+ ATPase PcdA is an early cell division protein that localizes to the future division site in a chromosome-independent manner before the cell splits into two equally sized daughter cells. PcdA forms a ring that constricts as the septum is being synthesized, following the migration of the FtsZ ring, and deletion of *pcdA* led to increased cellular area, misplacement of FtsZ, and defects in orthogonal plane selection. Mechanistically, PcdA interacts directly with unpolymerized FtsZ in an ATP-dependent fashion, and with FtsZ polymers. Additionally, PcdA interacts with DivIVA, a population of which we observe also localizes to the future cell division plane. The data are therefore consistent with a model (Fig. 4h) in which DivIVA marks the future cell division site in the two daughter cells as cell division proceeds in the parental cell, then recruits PcdA, which, in turn, recruits FtsZ. Consistent with this model, we observed that deletion of *divIVA* also results in an orthogonal cell plane selection defect. Thus, *S. aureus* joins a growing list of bacteria that employ a positive regulatory system to dictate correct placement of the cell division machinery ^43,44^.

While several key cell-division proteins such as FtsZ and DivIVA (interaction partners of PcdA) or chromosome segregation-associated factors (e.g., Noc, a member of the ParB superfamily) show a widespread or even pan-bacterial distribution, PcdA is unique in being limited to a lineage of coccoid Firmicutes. Our analysis indicates that PcdA is a late-emerging component that was derived from an McrB NTPase. This functional shift is puzzling because classical McrB NTPases function together with their restriction endonuclease partner in anti-viral immunity. How, then, did a conflict system give rise to a cell division protein? Given that PcdA is nested deeply within the radiation of McrBs from restriction systems, we hypothesize that certain McrB systems may also localize to cell division septa and protect sister cells from an invading virus by potentially restricting their DNA at that location. If so, PcdA could have emerged from such a precursor, with major structural changes and substitutions that converted its ancestral processive NTPase activity to a weak NTPase switch. Remarkably, we observe a parallel to this transformation of an McrB from an immunity factor to a cytoskeleton-associated protein in the animal CTTNBP2/ Nav2/Unc-53 lineage of AAA+ proteins. These proteins were derived from a bacterial McrB progenitor at the base of the animal lineage ^20^, where they diversified into versions that interact with F-actin as well as microtubules (eukaryotic cognates of FtsZ rings) in processes such as neuronal path-finding ^45,46^. Notably, CTTNBP2, which shows a parallel degeneration of the AAA+ NTPase domain as PcdA, interacts with tubulin to stabilize microtubules during the formation of dendritic spines ^46^, whereas PcdA interacts with the bacterial tubulin homolog FtsZ to direct FtsZ polymerization at the correct subcellular location.

Although key motifs required for nucleotide binding and hydrolysis are largely (Walker A and C-terminal AAA+-specific α-helical bundle) or partly (Walker B) eroded in PcdA and in its orthologs from certain coccoid Firmicutes, we discovered that PcdA weakly (but specifically) hydrolyzes ATP and GTP in vitro, suggesting that nucleotide hydrolysis may function as a switch to regulate PcdA function. Accordingly, we observed that further disrupting conserved residues in the AAA+ domain of PcdA resulted in a cell division defect in vivo. Furthermore, we found that nucleotide hydrolysis was required for PcdA multimerization and ATP was required for interaction with FtsZ. We also observed that PcdA correctly marked new cell division sites even in the absence of nucleoid, indicating that the PcdA ring is epigenetically inherited from the previous cell division event, and that the presence of the nucleoid does not primarily dictate selection of the next cell division event. Nonetheless, given the bias towards approximately orthogonal angles even in cells lacking PcdA, it is likely that not all angles are available for FtsZ polymerization and that the presence of the chromosome could restrict certain angles, likely via Noc inhibition of FtsZ polymerization over the chromosome. Since we did not detect an interaction between PcdA and Noc, the chromosome-restricted pathway is likely a parallel pathway to the positive regulation of FtsZ placement by PcdA.

A persistent question relates to the localization mechanism of DivIVA, which recruits PcdA to the correct division plane. DivIVA has been extensively studied in *B. subtilis*, where it is a structural protein that assembles into a platform to recruit other proteins during cell division and chromosome segregation during sporulation ^47^, but its function in *S. aureus* has remained mysterious ^37^. In *B. subtilis*, DivIVA preferentially binds to negatively curved membranes ^48,49^. Although *S. aureus* is considered spherical, it nonetheless slightly elongates prior to dividing ^9^, leading to the elaboration of “poles”. We propose that these differences in gross membrane curvature could drive the accumulation of DivIVA at those points, which coincides with a roughly orthogonal plane relative to the previous round of cell division (Fig. 4g), to recruit PcdA to that site.

Although deletion of *pcdA* did not result in an obvious growth defect in rich laboratory medium, we observed that it resulted in a virulence defect in a murine infection model. *Staphylococcus* abscesses in mouse models are characterized as persisting over time; following rupture, they release bacteria into the peritoneal cavity leading to new infection sites ^39^. When infected with the Δ*pcdA* strain, we observed that mice developed fewer abscesses. Further, although the Δ*pcdA* strain caused kidney lesions, they displayed an increased number of lesions that did not contain any bacteria compared to infection with the WT strain, which suggested that the Δ*pcdA* strain was more susceptible to immune clearance. We therefore propose that orthogonal cell division likely leads to the characteristic cluster-like growth of *S. aureus* and that this mode of growth may provide an advantage to evading the host immune system during infection. Given our observation that deletion of *pcdA* resulted in increased antibiotic sensitivity, we propose that PcdA represents a complementary antibiotic target to increase drug efficacy and promote clearance of staphylococcal infections that are otherwise recalcitrant to treatment by currently available antibiotics.

## ACKNOWLEDGMENTS

We thank S. Gottesman, G. Storz, S. Wickner, A. Khare, and T. Le for discussions, and P. Eswara and members of the K.S.R. laboratory for comments on the manuscript. Molecular graphics and analyses performed with UCSF Chimera, developed by the Resource for Biocomputing, Visualization, and Informatics at the University of California, San Francisco, with support from NIH P41-GM103311. This work was funded by the National Institutes of Health (NIH), the National Institute of Allergy and Infectious Diseases grant AI038897 (D.M) and 1R21AI156574 (J.L.C.), and the Intramural Research Program of the NIH, the National Cancer Institute, the Center for Cancer Research (K.S.R.), and the National Library of Medicine (L.A.). This work utilized the NIH HPC Biowulf computer cluster (V.A., L.A., C.T.).

## METHODS

### Bacterial strains, culture conditions, and plasmid construction

*Escherichia coli* strains were grown in Miller LB Broth (KD Medical) and LB agar. The medium was supplemented with 100 µg ml^-1^ spectinomycin, 100 µg ml^-1^ ampicillin, 50 µg ml^-1^ kanamycin, or 10 µg ml^-1^ chloramphenicol for plasmid maintenance, as required.

*Staphylococcus aureus* strains were grown in Tryptic Soy Broth (TSB) medium or modified M63 medium (13.6 g L^-1^ KH_2_PO_4_, 2 g L^-1^ (NH_4_)2SO_4_, 0.8 µM ferric citrate, 1 mM MgSO_4_; pH adjusted to 7 using KOH) supplemented with 0.3% glucose, 1X ACGU solution (Teknova, California, USA), 1X supplement EZ (Teknova), 0.1 ng L^-1^ biotin, and 2 ng L^-1^ nicotinamide. When required, medium was supplemented with 5 µg ml^-1^ erythromycin, 10 µ ml^-1^ chloramphenicol, or 1.5 µg ml^-1^ tetracycline. For growth on solid media, Tryptic Soy Agar (TSA) was used, supplemented with antibiotics at the above-indicated concentrations as needed.

To delete *pcdA*, upstream and downstream regions of *sausa300_2094* were amplified using primers 2094del-Gibson-F1 (tggatcccccgggctgcaggtgtaataacaatttaggagtgaatg) and 2094del-Gibson-R1 (tatatcattattctgctgtcatcttaatac), and 2094del-Gibson-F2 (gacagcagaataatgatatatggtagtttttgaaaag) and 2094del-Gibson-R2 (taccgggccccccctcgaggatttaatacactccttaaaattgtc), respectively. Fragments were cloned into pIMAY* digested with *EcoR*I and *Sal*I by Gibson assembly. The resulting plasmid was named pFRL134. For complementation of *pcdA* mutants, *sausa300_2094* and its promoter region were PCR-amplified using primers 2094-compl-F2 (atgaattctaattgtcgatagcgcg) and 2094-compl-R2 (atggatccaacgagtaatctatcataagctc) and cloned into pLL29 ^50^ using *BamH*I and *EcoR*I. The resulting plasmid was named pFRL112. For complementation with point mutations, site-directed mutagenesis was performed on pFRL112 using QuikChange Lighting Site-directed Mutagenesis kit (Agilent).

For localization of PcdA, *pcdA* gene and its promoter region was amplified using primers 2094_fwd (ttacccgtcttactgtcgggtaattgtcgatagcgcgtttg) and 2094_rev (cgctgcctccgtcatgtttgactttgactatac), and gene encoding superfolder GFP using primers 2094sGFP_fwd (caaacatgacggaggcagcggaatgagc) and 2094sGFP_rev (cctgcaggtcgactctagagtcatttgtagagctcatccatgc). The fragments were cloned into pLL29 digested with EcoRI and BamHI by Gibson assembly. The resulting plasmid was named pFRL126. For localization of point mutations, site-directed mutagenesis was performed on pFRL126 using QuikChange Lighting Site-directed Mutagenesis kit (Agilent). To obtain a pFRL126-derivative plasmid lacking the L54a *attP* site, pFRL126 was subjected to site-directed mutagenesis using the primers pPhi11-F (catgttgccaaaaatcgattatgtccagatctggaattaatgaggcattctaac) and pPhi11-R (gttagaatgcctcattaattccagatctggacataatcgatttttggcaacatg), originating the plasmid pFRL197.

For co-localization studies, *ezrA* and its promoter region were amplified using primers ezrA-mCherry-F (gaggccctttcgtcttcaagggcttgctgcttgtttctttaataatg) and ezrA-mCherry-R (ctcaccattccgctgcctccttgcttaataacttcttcttcaatatgtttag), and *mCherry* using primers GGSG-mCherry-F (ggaggcagcggaatggtgagcaagggcgaggagg) and GGSG-mCherry-R (ccctccggatccccgggtacttacttgtacagctcgtccatg). Both fragments were cloned into pCL55 ^51^ digested with EcoRI and KpnI by Gibson assembly. The resulting plasmid was named pFRL199. For localization of FtsZ-mCherry under an Cd2+-inducible promoter, *ftsZ* encoding region was amplified using primers Cd-ftsZ-F1 (aaggtcaatgtctgaacctgcaggctaggaggaaatttaaatgttag) and Cd-ftsZ-R1 (tccgctgcctccacgtcttgttcttcttgaac), and *mCherry* was amplified using primers Z-mCherry-F1 (agaacaagacgtggaggcagcggaatggtg) and Z-mCherry-R1 (tatgcattagaataggcgcgcctgttacttgtacagctcgtccatgc), and fragments were cloned into pJB67 ^52^ digested with SalI and EcoRI. The resulting plasmid was named pFRL221. For complementation of Tn::*scpB*, scpB gene and its promoter were amplified using primers 1444-pLL29-F (ttacccgtcttactgtcggggtataacgcatctctatctttag) and 1444-pLL29-R (cctgcaggtcgactctagaggccttacgtcttgaagtataac), and cloned into pLL29 digested with EcoRI and BamHI using Gibson assembly. The resulting plasmid was named pFRL110.

For overexpression and purification of His_6_-PcdA, *pcdA* encoding sequence was amplified using primers 2094his-F (ctggtgccgcgcggcagccatatgacagcagaaacgaattatttttg) and 2094his-R (gtcgacggagctcgaattcgttagtcatgtttgactttgac), and cloned into pET28a digested with NdeI and BamHI using Gibson assembly. The resulting plasmid was named pFRL132. For purification of untagged PcdA, *pcdA* was amplified using primers pcdA-sumo-F (ctcacagagaacagattggtggtatgacagcagaaacgaattatttttg) and pcdA-sumo-R (tcgggctttgttagcagccgttagtcatgtttgactttgac) and cloned into pTB146 digested with BamHI and SapI using Gibson assembly. The resulting plasmid was called pFRL159. For purification of PcdA^T430A^, site-directed mutagenesis was performed on pFRL159 resulting in the plasmid pFRL194.

For bacterial two-hybrid assays, pUT18 was linearized with primers pUT18-Gibson-F (gtaccgagctcgaattcagcc) and BTH-Gibson-R (catagctgtttcctgtgtgaaattg), and assembled with *pcdA* amplified with primers pcdA-BTH-fwd (tcacacaggaaacagctatgacagcagaaacgaattatttttg) and pcdA-T18-rev (gctgaattcgagctcggtacgtcatgtttgactttgac), resulting in plasmid pFRL161. For cloning into pKNT25, *pcdA* (amplified with primers pcdA-BTH-fwd and pcdA-T25-rev [attgaattcgagctcggtacgtcatgtttgactttgac]), *zapA* (amplified with primers zapA-BTH-fwd [tcacacaggaaacagctatgacacagtttaaaaacaaggtaaatgtatcaattaatgatc] and zapA-T25-rev [attgaattcgagctcggtacttgctcacgctgctgcaatttg]), *ezrA* (amplified with primers ezrA-BTH-fwd [tcacacaggaaacagctatggtgttatatatcattttggcaataattg] and ezrA-T25-rev [attgaattcgagctcggtacttgcttaataacttcttcttcaatatg]), *noc* (amplified with primers noc-BTH-fwd [tcacacaggaaacagctatgaaaaaacctttttcaaaattatttgg] and noc-T25-rev [attgaattcgagctcggtacacgtttatatattcgaatttttatttc], *spo0J* (amplified with primers spo0J-BTH-fwd [tcacacaggaaacagctatgagtgaattgtcaaaaagtgaag] and spo0J-T25-rev [attgaattcgagctcggtactttaccatacctacgatttaattg]), and *divIVA* (amplified with primers divIVA-BTH-fwd [tcacacaggaaacagctatgccttttacaccaaatgaaattaag] and divIVA-T25-rev [attgaattcgagctcggtaccttcttagttgtttctgaatc]) were cloned into pKNT25 linearized with primers pKNT25-Gibson-F (gtaccgagctcgaattcaatgacc) and BTH-Gibson-R using Gibson assembly. Resulting plasmids were pFRL168 (*pcdA*), pFRL170 (*zapA*), pFRL171 (*ezrA*), pFRL172 (*noc*), pFRL173 (*spo0J*), and pFRL174 (*divIVA*). For variants of the *pcdA*, site-directed mutagenesis was performed on plasmids pFRL161 and pFRL168.

### Strain construction

*Staphylococcus aureus* strains used in this study are derivates of JE2 ^15^, a plasmid-less derivative of the methicillin-resistant *S. aureus* (MRSA) USA300 lineage. For deletion of *sausa300_2094* (*pcdA*), plasmid pFRL134 was transformed into *S. aureus* RN4220 ^53^ and maintained at 30⁰C until transduced into *S. aureus* JE2 strain using φ85. Allelic exchange was carried out as described previously ^54^. Briefly, single crossover event was selected by growth in TSB medium at 37⁰C in the presence of 10 μg ml^-1^ chloramphenicol. After growing overnight in the absence of chloramphenicol, double crossover and loss of the plasmid was selected by plating on TSA in the presence of 40 mM *para*-chlorophenylalanine (PCPA). Several clones were tested and deletion of *pcdA* was confirmed by PCR. One clone carrying the deletion of *pcdA* was selected and named FRL60. For genomic integration into the φ11 attB site (for complementation and expression of sGFP fusions), plasmids pFRL112, pFRL126, or derivatives were transformed by electroporation into *S. aureus* RN4220 bearing plasmid pLL2757, which encodes the integrase for φ11. Transformants were selected by growing on TSA plates containing 1.5 μg ml^-1^ tetracycline. Integration of the plasmid in the φ11 attB site was confirmed by PCR and then transduced into strain JE2-derived using φ85.

For co-localization of PcdA and EzrA, strain carrying plasmid pFRL197 was transduced with lysates of RN4220 bearing pFRL199 inserted in L54a *attP* site. Clones were selected on TSA plates containing 1.5 μg ml^-1^ tetracycline and 10 μg ml^-1^ chloramphenicol. Genomic DNA of the transductants was isolated and correct insertion of both plasmids was confirmed by PCR. For co-localization of PcdA and FtsZ, strain carrying plasmid pFRL197 was transduced with lysates of RN4220 bearing pFRL221. Clones were selected on TSA plates containing containing 1.5 μg ml^-1^ tetracycline and 5 μg ml^-1^ erythromycin. For localization of FtsZ-mCherry, pFRL221 was transduced into JE2 wild type background or Δ*pcdA* (FRL60 strain).

### Genetic selection for mutants defective in cell division

Individual mutants in the Nebraska Transposon Mutant Library (NTML) ^15^ were grown overnight in 700 μl modified M63 medium in deep 96-well plates at 37 ⁰C. Overnight cultures were then diluted 1:100 in 150 μl modified M63 medium alone or supplemented with 100 ng ml^-1^ PC190723^17^ in 96-well plates. Cultures were then incubated at 37 ⁰C, with shaking, and optical density at 600 nm was continuously monitored. Mutants whose growth curves were attenuated in the presence of PC190723 compared to WT were validated by growing them in 96-well plates as indicated above in the absence or presence of PC190723 and monitoring their growth kinetics.

### Analysis of protein expression by immunoblot

Strains were grown in modified M63 medium until reaching mid-exponential phase. OD_600_ was adjusted for all strains to 0.4 and cells from 4 mL culture were pelleted by centrifugation (7,500 x *g*, 10 min). Pellet was then resuspended in Lysis Buffer (50 mM Tris-HCl pH 7.5, 150 mM NaCl, 5 mM EDTA, 0.1 mg mL-1 lysostaphin, 0.1 mg mL-1 lysozyme) and incubated at 37⁰C for 5 min. Samples were then diluted by adding 1:2 volume Wash Buffer (50 mM Tris-HCl pH 7.5, 150 mM NaCl, 5 mM EDTA) and loading buffer, and incubated at 95⁰C for 5 min. Equivalent volume of all samples were loaded in a SDS-PAGE gel, and then transferred to a PVDF membrane. Blocking of the membrane was carried out by incubating in blocking solution for 1 h at room temperature (5% Bio-Rad Blotting-Grade Blocker in TSB-T buffer). Membrane was then incubated with anti-PcdA polyclonal antibody diluted 1:15,000 in blocking solution for 2 h at room temperature.

Membrane was then washed three times with TBS-T and incubated with secondary antibody (Goat Anti-Rabbit IgG StarBright Blue 700 by Bio-Rad) diluted 1:3,000 in blocking solution for 45 min. After washing three times with TBS-T, signal was visualized using a ChemiDoc Imaging System (Bio-Rad).

### Purification of His_6_-PcdA, untagged PcdA, and untagged FtsZ

Rabbit anti-PcdA antiserum was produced against purified His_6_-PcdA. Briefly, overnight cultures of *E. coli* BL21(DE3) carrying pFRL132, grown in LB medium containing 50 μg ml^-1^ kanamycin, were diluted 1:50 into 500 ml fresh LB medium containing kanamycin and grown until OD_600_ reached 0.4. At this time, protein expression was induced by adding isopropyl β-d-1-thiogalactopyranoside (IPTG) at 0.5 mM final concentration, and cultures were incubated at 37⁰C, shaking at 250 rpm for 3 h. Cells were harvested by centrifugation and resuspended in Lysis Buffer (100 mM NaH_2_PO_4_, 10 mM Tris, 8 M urea, pH 8.0). Cell disruption was carried out using a French pressure cell at ca. 20,000 psi. Cell lysate was cleared by centrifugation at 100,000 × *g* for 30 min at 4 ⁰C and then incubated with Ni^2+^-NTA agarose beads (Qiagen) for 30 min at 4⁰C. The beads were then transferred to a column, washed with Lysis Buffer, and eluted with Elution Buffer (100 mM NaH_2_PO_4_, 10 mM Tris, 8 M urea, pH 4.5). Protein purity was assessed by Coomassie-stained SDS-PAGE and purified material was used to immunize rabbits (Labcorp, USA).

To purify untagged PcdA or PcdA^T430A^ variant, *E. coli* BL21(DE3) carrying pFRL159 or pFRL194, respectively, was grown in LB medium containing 50 μg ml^-1^ ampicillin and 1% glucose overnight at 37 ⁰C at 250 rpm. Overnight cultures were then diluted 1:50 into 1 L fresh LB/ampicillin and incubated at 37 ⁰C at 250 rpm. When OD_600_ reached 0.4, protein expression was induced by adding 0.5 mM IPTG (final concentration) and cultures were incubated at 37 ⁰C, 250 rpm for 4 h. Cells were harvested by centrifugation, resuspended in Buffer A (50 mM Tris-HCl pH 8.0, 150 mM KCl, 10% glycerol), and disrupted using a French pressure cell at ca. 20,000 psi. Cell lysate was cleared by centrifugation at 100,000 × *g* for 30 min at 4 ⁰C. The supernatant was then incubated with Ni^2+^-NTA agarose beads (Qiagen) for 30 min at 4 ⁰C, applied to a column, and washed with three column volumes of Wash Buffer (Buffer A containing 30 mM imidazole). Protein was then eluted using Elution buffer (Buffer A containing 300 mM imidazole), after which the imidazole was promptly removed using a PD-10 desalting column (Cytiva). The resulting protein solution was incubated overnight at 4 ⁰C in the presence of His_6_-Ulp1 protease (100:1 ratio) in Buffer A containing 1 mM DTT. The cleaved His_6_-SUMO tag and His_6_-Ulp1 protease were then removed by incubation with Ni^2+^-NTA agarose beads as described earlier. The flowthrough fraction containing untagged PcdA was confirmed by SDS-PAGE, quantified by Bradford assay, and stored at −80 ⁰C until further use. S. aureus FtsZ was purified as previously described ^55^.

### Microscopy

Overnight cultures of *S. aureus* in modified M63 medium, containing 1.5 μg ml^-1^ tetracycline or 10 μg ml^-1^ chloramphenicol, if necessary, were diluted 1:100 into fresh medium and grown to mid-logarithmic phase. 1 ml culture was then washed with complete M63 and resuspended in ∼100 μl PBS containing 10 μg ml-1 fluorescent dye FM4-64 and/or 2 μg ml^-1^ DAPI to visualize membranes and nucleoid, respectively. 5 μl was spotted on a glass bottom culture dish (Mattek) and covered with a 1% agarose pad made with PBS. Cells were imaged using a DeltaVision Core microscope system (Applied Precision/GE Healthcare) equipped with a Photometrics CoolSnap HQ2 camera. Data were deconvolved using SoftWorx software. Cellular area was measured using Fiji software by manually tracing the cell contour in FM4-64-stained cells.

Analysis of consecutive planes of cell division was performed as described previously ^6^. Briefly, *S. aureus* strains were grown in modified M63 medium to mid-logarithmic phase and then WGA-488 (Invitrogen) was added to 1 μg ml^-1^ final concentration and incubated for 5 min at room temperature. Cells were then collected by centrifugation, resuspended in modified M63 pre-warmed at 37 ⁰C, and incubated in darkness at 37 ⁰C at 250 rpm for 40 min to ensure one round of cell division. 1 ml culture was then pelleted and cells were resuspended in 100 μl PBS containing 10 μg ml^-1^ fluorescent dye FM4-64. Cells were imaged as indicated taking Z-stacks of 20 slides with a spacing of 0.15 μm. Angle between consecutive planes was measured by tracing the borders of WGA-488 staining and septum labeled with membrane dye.

### Quantification of NTP hydrolysis

Purified PcdA or PcdA^T430A^ were prepared at 2.5 μM final concentration in reaction buffer (50 mM Tris-HCl at pH 8.0, 150 mM KCl, 5 mM MgCl_2_). Reactions were initiated by addition of ATP, GTP, CTP, or UTP at final concentrations of 0.25, 0.5, 1, 2, and 4 mM. Reactions were then incubated at 37 ⁰C and stopped after 30 min by adding an equal volume of 20 mM EDTA. Reactions were transferred to a 96-well plate and the amount of inorganic phosphate released from the hydrolysis of NTP was determined using Phosphate Assay Kit PiColorLock (Abcam) according to protocol by the manufacturer, using a standard curve of known concentrations of inorganic phosphate.

### Protein-Protein interaction assays

For *in vitro* assays, purified *S. aureus* FtsZ (30 μM) was incubated in reaction buffer (20 mM HEPES at pH 7.5, 140 mM KCl, 5 mM MgCl_2_) in the absence or presence of PcdA (10 μM) and 2 mM non-hydrolysable GTP analog GMPPCP. Reaction was incubated for 10 min at 37 ⁰C and then centrifuged for 30 min at 129,000 × *g*. Supernatant and pellet fractions were prepared at equivalent volumes and analyzed by SDS-PAGE. Coomassie staining and protein bands were quantified using ImageJ software.

For retention assays, FtsZ (30 μM) and PcdA (10 μM) were incubated in reaction buffer in the absence or presence of 2 mM ATP. After incubation for 15 min at room temperature, reactions were applied to prewashed 100 kDa retention filter (Pall Life Sciences) and centrifuged at 21,000 × *g* for 10 min. Flowthrough and retained fraction were prepared at equivalent volumes and analyzed by SDS-PAGE. Quantification of protein bands was done using ImageJ after Coomassie staining.

For bacterial two-hybrid assays, pUT18 and pKNT25 derived plasmids were transformed into *E. coli* strain BTH101 (*cyaA*-), and transformants were selected on LB plates containing 100 μg ml^-1^ ampicillin, 50 μg ml^-1^ kanamycin, and 1% glucose at 30 ⁰C. For each combination of plasmids, 3 colonies were pooled together and grown shaking overnight at 30 ⁰C in LB medium containing ampicillin, kanamycin, and 1 mM IPTG. The next day, cells were diluted 1:10 into fresh LB medium and OD_600_ was measured for each culture. 100 μl diluted cell solution was lysed by adding 10 μl Lysis Buffer (1 mg ml^-1^ lysozyme in 1X BugBuster buffer (Sigma) and incubated at room temperature for 15 min. Interaction between fusion proteins was quantified as β-galactosidase activity. To measure it, lysed cells were diluted 1:1 with Z Buffer (62 mM Na_2_HPO_4_, 45 mM NaH_2_PO_4_, 10 mM KCl, 1 mM MgSO_4_, 50 mM β-mercaptoethanol) and reaction was started by adding ortho-Nitrophenyl-β-galactoside (ONPG) to 2 mM final concentration. Hydrolysis of ONPG was monitored by measuring OD_420_ every 5 seconds for 30 min using a microplate reader (Tecan). β-galactosidase activity was determined from the linear range of the curves obtained when OD_420_ plotted against time.

### Mouse renal abscess studies

BALB/c mice, 6-8 weeks of age, were obtained from Jackson Laboratory and infected with the wild type JE2 strain (WT) or the isogenic Δ*pcdA* variant in groups of up to 10 animals. Inocula for infection were prepared by growing bacterial cultures in tryptic soy broth to A600 of 0.42. Cells were sedimented, washed once with PBS, and then suspended in PBS. Animals were anesthetized with a cocktail of ketamine-xylazine (50 to 65 and 3 to 6 mg/kg) and injected into the periorbital venous plexus with a 100-µl suspension containing ∼1 x 10^7^ colony forming units (CFU). The size of inocula was verified by platting bacterial suspensions on tryptic soy agar at 37 °C followed by enumeration of CFU. Following infection, mice were monitored for clinical signs of disease and body weight was recorded daily for a total of 15 days. On days 5 and 15 post-infection, animals were euthanized by carbon dioxide inhalation. Kidneys were collected and surface abscesses were enumerated. Kidneys were then fixed in 10% formalin for 24 h at room temperature, embedded in paraffin, thin sectioned, stained with hematoxylin-eosin, and inspected by light microscopy to enumerate deep-seated abscess lesions.

### Sequence Analysis

Sequence similarity searches were performed using the PSI-BLAST program with a profile-inclusion threshold set at an e-value of 0.01 ^56^. The searches were conducted against the NCBI non-redundant (nr) database or the same database clustered down to 50% sequence identity using the MMseqs program, or a curated database of 7423 representative genomes from across the tree of life. Profile-profile searches were performed with the HHpred program ^57,58^. Multiple sequence alignments (MSAs) were constructed using the FAMSA and MAFFT programs ^59,60^. Sequence logos were constructed using these alignments with the ggseqlogo library for the R language ^61^.

### Structure Analysis

PDB coordinates of structures were retrieved from the Protein Data Bank and structures were rendered, compared, and superimposed using the Mol* program ^62^ and UCSF Chimera ^63^. Structural models were generated using the AlphaFold2 and AlphaFold-Multimer programs ^30,64^. Multiple alignments of related sequences (>30% similarity) were used to initiate HHpred searches for the step of identifying templates to be used by the neural networks deployed by these programs.

### Comparative Genomics and Phylogenetic Analysis

Clustering of protein sequences and the subsequent assignment of sequences to distinct families was performed by the MMSEQS program, adjusting the length of aligned regions and bit-score density threshold empirically. Phylogenetic analysis was performed using the maximum-likelihood method with the IQTree program and multiple protein substitution models such as Dayhoff, Poisson, and JTTMutDC. The FigTree program (http://tree.bio.ed.ac.uk/software/figtree/) was used to render phylogenetic trees. Gene neighborhoods were extracted through custom PERL scripts from genomes retrieved from the NCBI Genome database.

## Ethics Statement

Animal research was performed in accordance with institutional guidelines following experimental protocol review, approval, and supervision by the Institutional Animal Care and Use Committee at The University of Chicago. Experiments with *S. aureus* were performed in Biosafety Level 2 containment.

## SUPPLEMENTAL DATA

**Figure S1.**
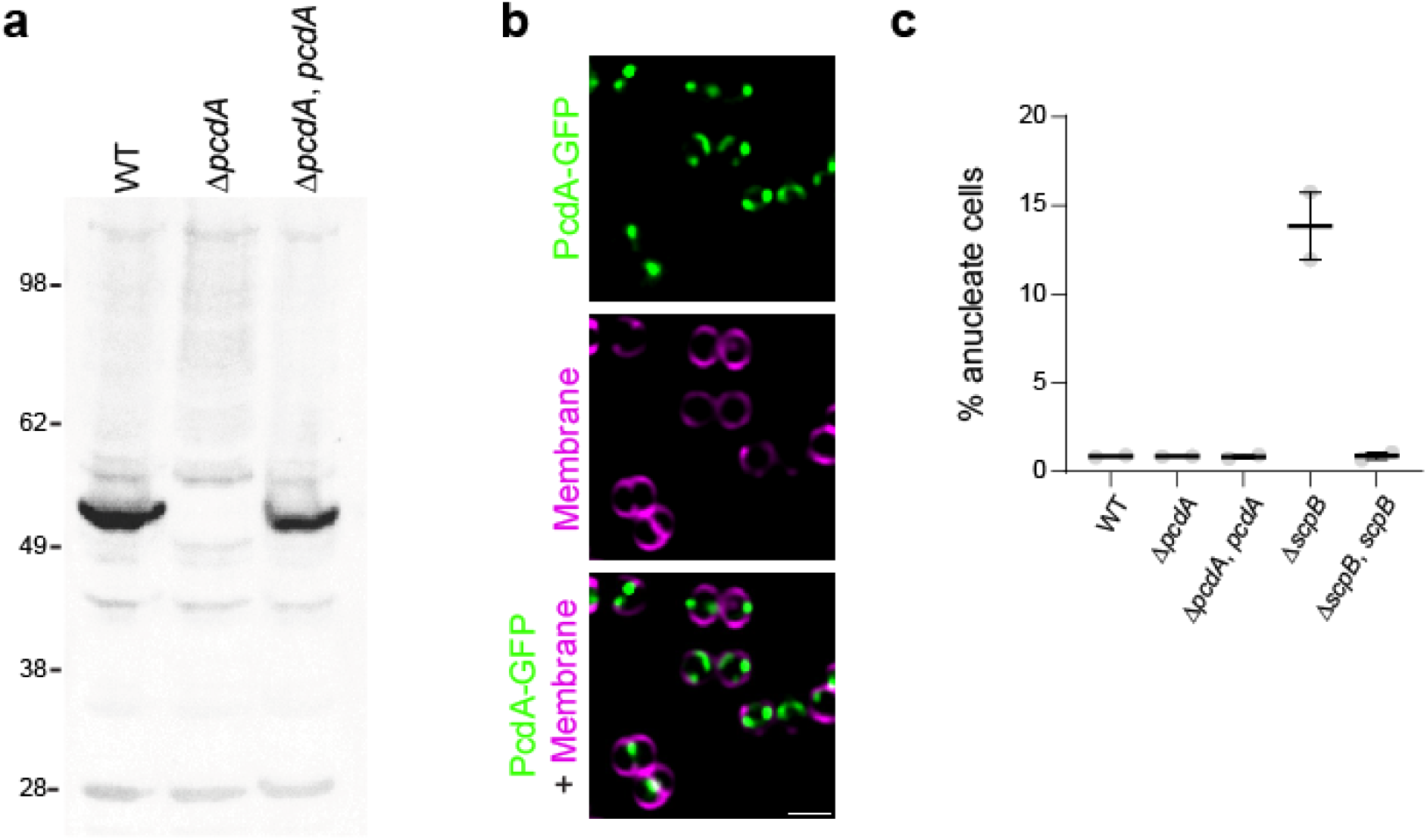
(a) Immunoblot using polyclonal antibodies against PcdA using extracts from WT, Δ*pcdA*, and Δ*pcdA* complemented at an ectopic chromosomal locus with *pcdA* strains. Predicted molecular weight for PcdA is ∼53 kDa. Strains: JE2, FRL60, and FRL62. (b) Larger field of view showing subcellular localization of PcdA-sGFP in WT strain. First row: PcdA-sGFP (green); second row: membrane stained with FM4-64 (magenta); third row: overlay of PcdA-sGFP and membrane. Scale bar: 1 μm. Strain FRL28. (c) Graph showing percentage of anucleate cells for WT, Δ*pcdA*, complemented Δ*pcdA*, Δ*scpB*, and complemented Δ*scpB*. Data points are from two independent replicates where > 1,000 cells were analyzed for each strain; bars represent averages; errors: SEM. Strains: FRL60, FRL62, NE1085, and FRL12.

**Figure S2.**
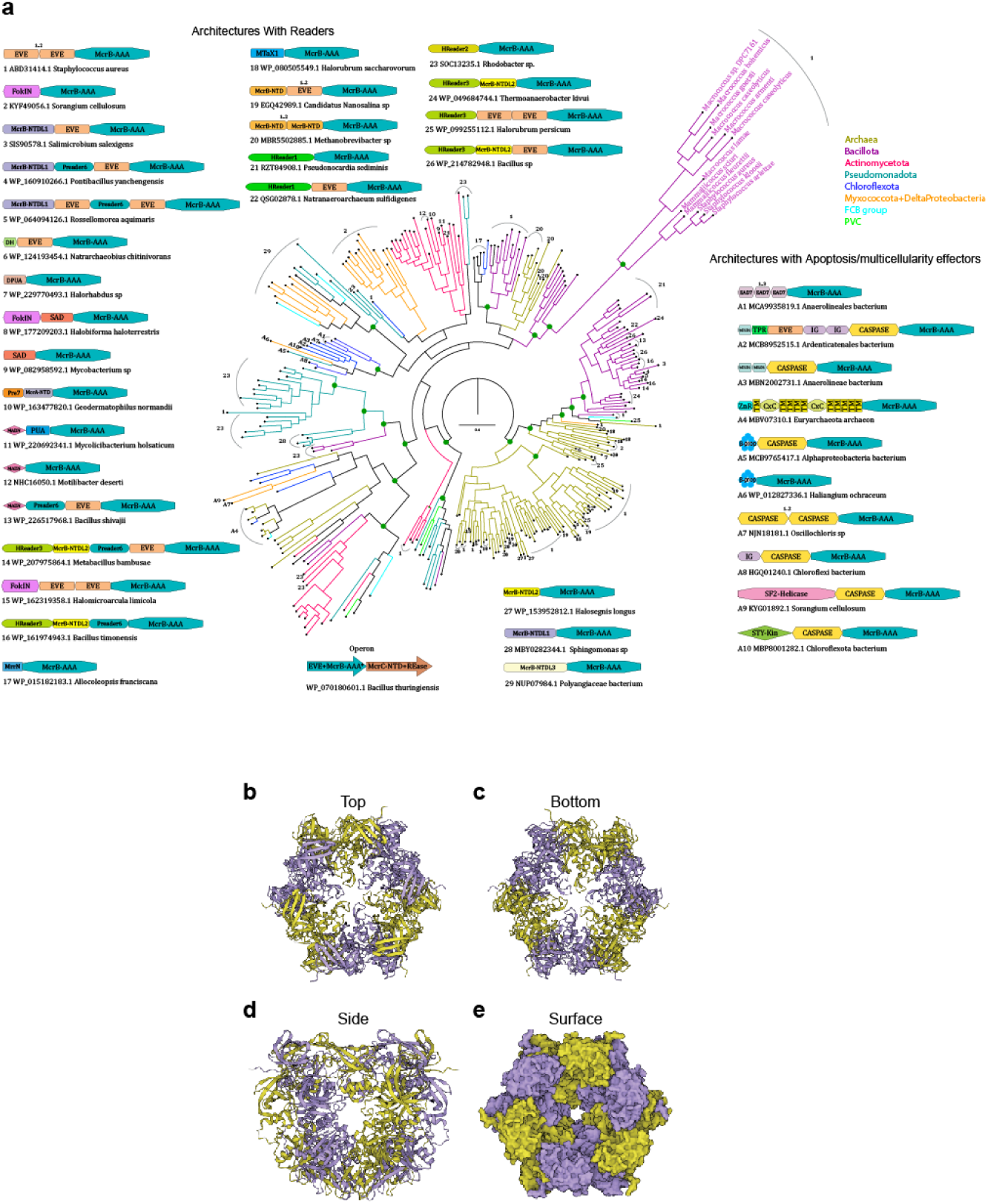
Phylogenetic tree of McrB AAA+ ATPase from a Multiple Sequence Alignment with a curated set of proteins. (a) The clades are colored by taxonomy as shown in the legend. Organism names are shown for the PcdA branch. Representative architectures and operon are shown with the accession and organism name below them. The arrows denote the genes in the operon with the “*” denoting the accession shown below it. The architectures are numbered, and the numbers are placed in the branches where they are found. Domains with variability in the number of tandem repeats are shown with a 1..2 or 1..3 above them. Abbreviations: FokIN – FokI-N-terminal-domain like, MAD-NTDL - MAD-NTD-like, DH - DpnI-HTH, DPUA - DCD-PUA, Pre7 - Prereader7, and MADN - MAD-NTD. The tree was generated using IQtree with the Dayhoff amino-acid exchange rate matrix which is empirically determined as one of the best fits. Key branches with boot strap support greater than 90% are shown with a green dot. (b-e) The predicted structure of a PcdA hexamer showing views along (b-c) the hexamer axis, (d) a side view perpendicular the axis, and (e) a surface view revealing a potential binding cavity. PcdA structures were generated using AF2.

**Figure S3.**
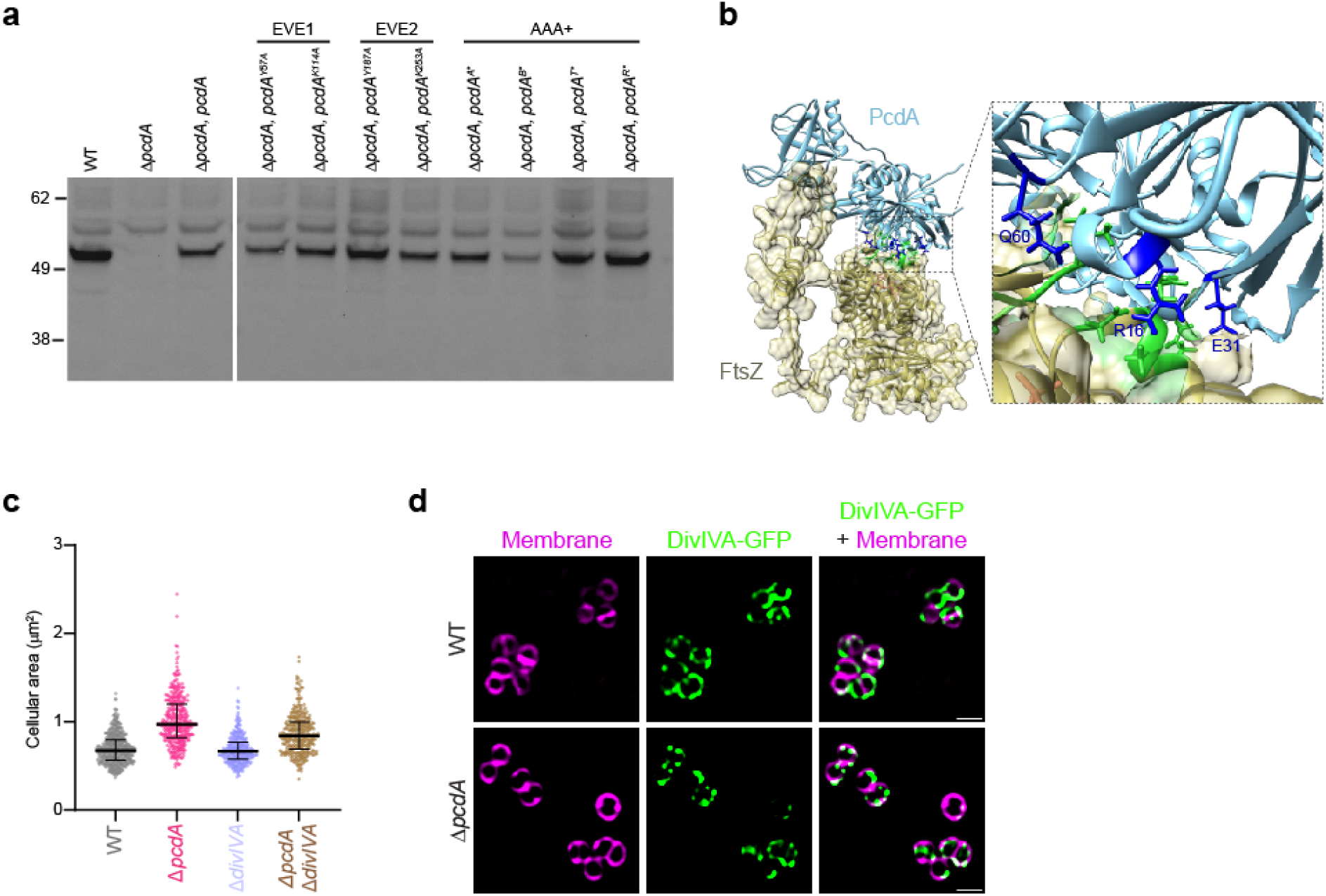
(a) Immunoblot using polyclonal antibodies against PcdA against cell extracts of WT, Δ*pcdA*, complemented Δ*pcdA*, or Δ*pcdA* complemented with indicated allele of *pcdA*. Strains: JE2, FRL60, FRL14, FRL34 – 41. (b) Protein complex model of a monomeric FtsZ (cyan) and PcdA (gold) predicted by AlphaFold-Multimer. A close-up section of the predicted interphase indicating residues R16, E31, and Q60 on PcdA may mediate the interaction with FtsZ. (c) Cellular area (μm^2^) of the indicated strains (n > 300 cells). Bars indicate median; whiskers indicate interquartile range. Strains: JE2, FRL60, FRL98, and FRL96. (d) Subcellular localization of DivIVA-sGFP in the WT and Δ*pcdA* strains. First column: membranes stained with FM4-64 (magenta); second column: DivIVA-sGFP (green); third column: overlay of membrane and DivIVA-sGFP. Scale bar: 1 μm. Strains: FRL113 and FRL114.

**Figure S4.**
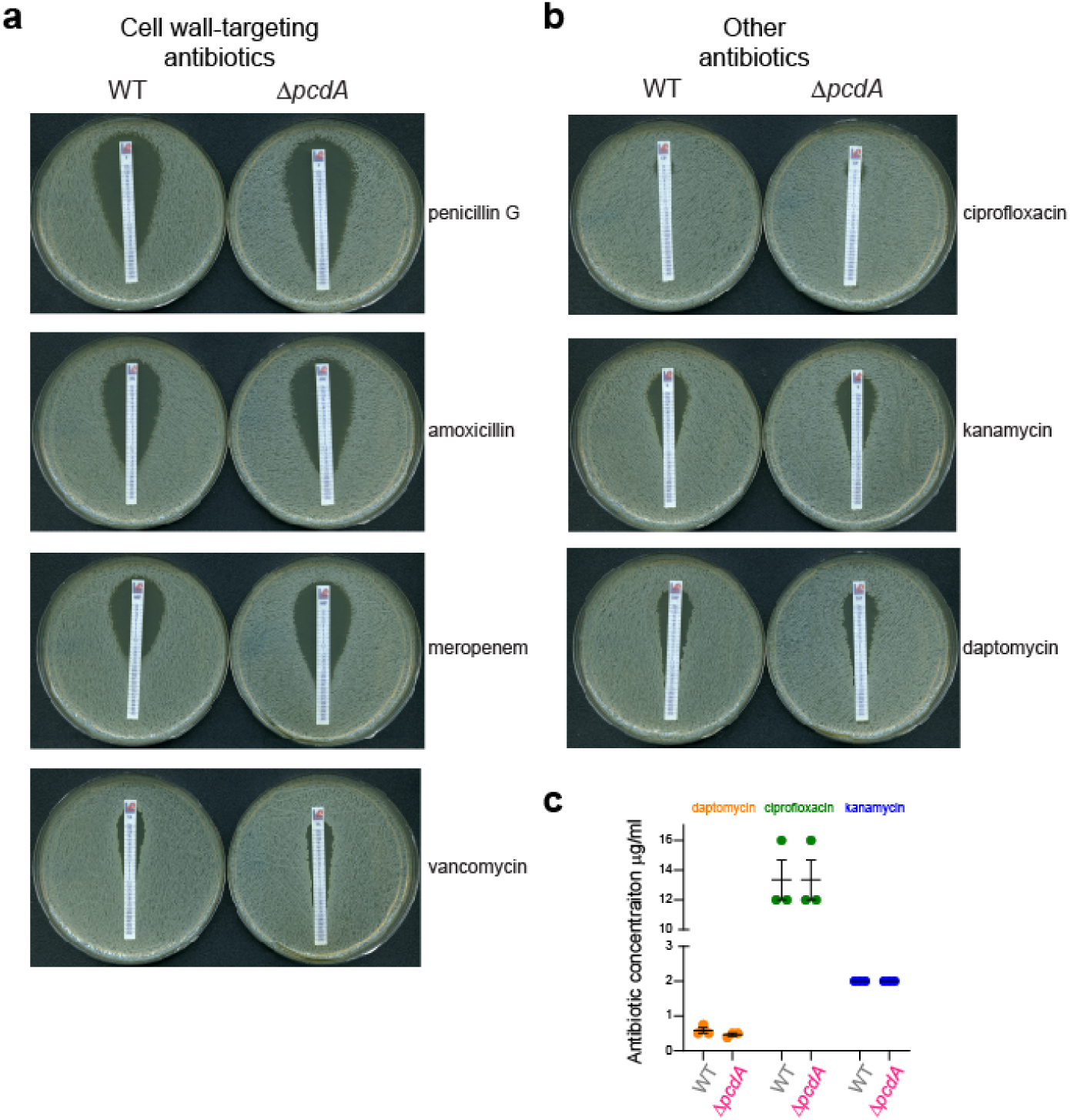
(a) Representative images of plates for MIC determination of cell wall-targeting antibiotics for the indicated strains. MIC was indicated by the intersection of the inhibition ellipse with the MIC strip. (b) Representative images of plates for MIC determination of antibiotics targeting other cellular process such as DNA metabolism (ciprofloxacin), protein synthesis (kanamycin), or cytoplasmic membrane (daptomycin). (c) MICs for JE2 wild type and Δ*pcdA* for the indicated antibiotics. Strains: JE2 and FRL60.

